# pGlycoQuant with a deep residual network for precise and minuscule-missing-value quantitative glycoproteomics enabling the functional exploration of site-specific glycosylation

**DOI:** 10.1101/2021.11.15.468561

**Authors:** Weiqian Cao, Siyuan Kong, Wenfeng Zeng, Pengyun Gong, Biyun Jiang, Xinhang Hou, Yang Zhang, Huanhuan Zhao, Mingqi Liu, Xihua Qiao, Mengxi Wu, Guoquan Yan, Chao Liu, Pengyuan Yang

**Affiliations:** Shanghai Fifth People’s Hospital, Fudan University, and Shanghai Key Laboratory of Medical Epigenetics, International Co-laboratory of Medical Epigenetics and Metabolism (Ministry of Science and Technology), Institutes of Biomedical Sciences, Fudan University, Shanghai, China; Key Lab of Intelligent Information Processing of Chinese Academy of Sciences (CAS), Institute of Computing Technology, CAS, Beijing, China; Beijing Advanced Innovation Center for Big Data-Based Precision Medicine, School of Engineering Medicine, Beihang University, Beijing, 100191, China; and Key Laboratory of Big Data-Based Precision Medicine (Beihang University), Ministry of Industry and Information Technology; NHC Key Laboratory of Glycoconjugates Research, Fudan University, Shanghai, China; Proteomics and Signal Transduction, Max Planck Institute of Biochemistry, Martinsried, Germany

## Abstract

Interpreting large-scale glycoproteomic data for intact glycopeptide identification has been tremendously advanced by software tools. However, software tools for quantitative analysis of intact glycopeptides remain lagging behind, which greatly hinders exploring the differential expression and functions of site-specific glycosylation in organisms. Here, we report pGlycoQuant, a generic software tool for accurate and convenient quantitative intact glycopeptide analysis, supporting both primary and tandem mass spectrometry quantitation for multiple quantitative strategies. pGlycoQuant enables intact glycopeptide quantitation with very low missing values via a deep residual network, thus greatly expanding the quantitative function of several powerful search engines, currently including pGlyco 2.0, pGlyco3, Byonic and MSFragger-Glyco. The pGlycoQuant-based site-specific N-glycoproteomic study conducted here quantifies 6435 intact N-glycopeptides in three hepatocellular carcinoma cell lines with different metastatic potentials and, together with in vitro molecular biology experiments, illustrates core fucosylation at site 979 of the L1 cell adhesion molecule (L1CAM) as a potential regulator of HCC metastasis. pGlycoQuant is freely available at https://github.com/expellir-arma/pGlycoQuant/releases/. We have demonstrated pGlycoQuant to be a powerful tool for the quantitative analysis of site-specific glycosylation and the exploration of potential glycosylation-related biomarker candidates, and we expect further applications in glycoproteomic studies.

## Introduction

Protein glycosylation has long been known as a heterogeneous posttranslational modification (PTM) that increases protein diversity and exerts a profound effect on various biological processes^1-4^. With great strides made in mass spectrometry (MS)-based analytical methods and interpretation software tools^5-10^, the identification of intact glycopeptides on a proteome-wide scale is no longer a serious obstacle to glycosylation analysis^11^.

However, the reliable and global quantitative analysis of intact glycopeptides remains a challenging barrier due to the lack of efficient software tools for the quantitative interpretation of proteome-scale intact glycopeptide mass spectrometry data^6, 12^. Although primary and tandem mass spectrometry (MS1/MS2)-based quantitative strategies, such as label-free, isotope chemical labeling and isotope metabolic labeling approaches, have been accepted as gold standard methods for proteomics quantitative analysis^13-15^, the wide application of these strategies in large-scale quantitative intact glycoproteomic studies has been clearly impeded by the lack of mature software tools for quantitative data processing. Among the few software tools available for intact glycopeptide quantitation^16^, almost all suffer from impaired accuracy of quantitation and a large amount of quantitative missing values caused by inappropriate quantitative data processing procedures^6, 12^.

A targeted mass spectrometry signal is easily affected by interference from nearby signals or noise, and its morphological characteristics cannot be entirely remained, resulting in impaired accuracy or missing values^17^. Deep learning-based algorithms have led to very good performance on a variety of subjects^18, 19^. Among them, the deep residual neural network (ResNet) introduced by He et al.^20^ has been accepted as an effective method for training computational vision object detection models that can represent much more complex functions than were previously practically feasible^21-23^. The main benefit of ResNet is that an image or matrix could be transformed to a well-trained vector that shows excellent performance in learning patterns from complex data and in matching two matrices^24-26^.

Here, we present pGlycoQuant, a dedicated software tool for large-scale and global quantitative glycoproteomics that applies the ResNet deep learning to processing glycopeptide quantitative evidence between or within MS runs. We applied pGlycoQuant to state-of-the-art glycopeptide quantification analysis and comparison with other quantitation software tools, including MSFragger-Glyco and Byologic™. pGlycoQuant reports 1/10,000 missing values for glycopeptide quantification with match-between-run analysis, which is two magnitudes less than that of other quantitative software tools. The current version of pGlycoQuant supports both primary and tandem mass spectrometry quantitation for multiple quantitative strategies, including label-free, chemical labeling and metabolic labeling approaches, and is compatible with identification results from several widely used search engines, including the Byonic^27^, MSFragger-Glyco^8^, Open-pFind^28^ and pGlyco series^7, 29^ engines. Furthermore, a pGlycoQuant-based site-specific N-glycoproteomic study quantified 6435 intact N-glycopeptides in three hepatocellular carcinoma (HCC) cell lines with different metastatic potentials and, together with in vitro molecular biology experiments, identified core fucosylation at site 979 of the L1 cell adhesion molecule (L1CAM) as a potential regulator of HCC metastasis.

## Results

### 1. Development and optimization of pGlycoQuant

The rapid development of software tools for large-scale glycoproteomic data interpretation has greatly facilitated intact glycopeptide identification^6^. However, only a few of them can be used for glycopeptide quantitation^30, 31^, and there is no mature software for quantitative data processing, mostly because of impaired accuracy and large numbers of quantitative missing values (Figure 1a). Efficient tools for the comprehensive and accurate quantitative interpretation of proteome-scale intact glycopeptide mass spectrometry data are lagging behind and greatly needed. We developed pGlycoQuant, a dedicated software tool, for large-scale and global quantitative glycoproteomics. pGlycoQuant supports both primary and tandem mass spectrometry quantitation for multiple quantitative strategies, including label-free, chemical labeling and metabolic labeling approaches, and is compatible with several widely used search engines, including Byonic^27^, MSFragger-Glyco^8^, Open-pFind^28^ and pGlyco series^7, 29^ (Figure 1b). pGlycoQuant consists of three steps: first reading the identification results from searching engines, then extracting the quantitation signals, and finally processing the quantitation results (Supplementary Figure 1). The whole workflows are described in the Online Methods.

**Figure 1.**
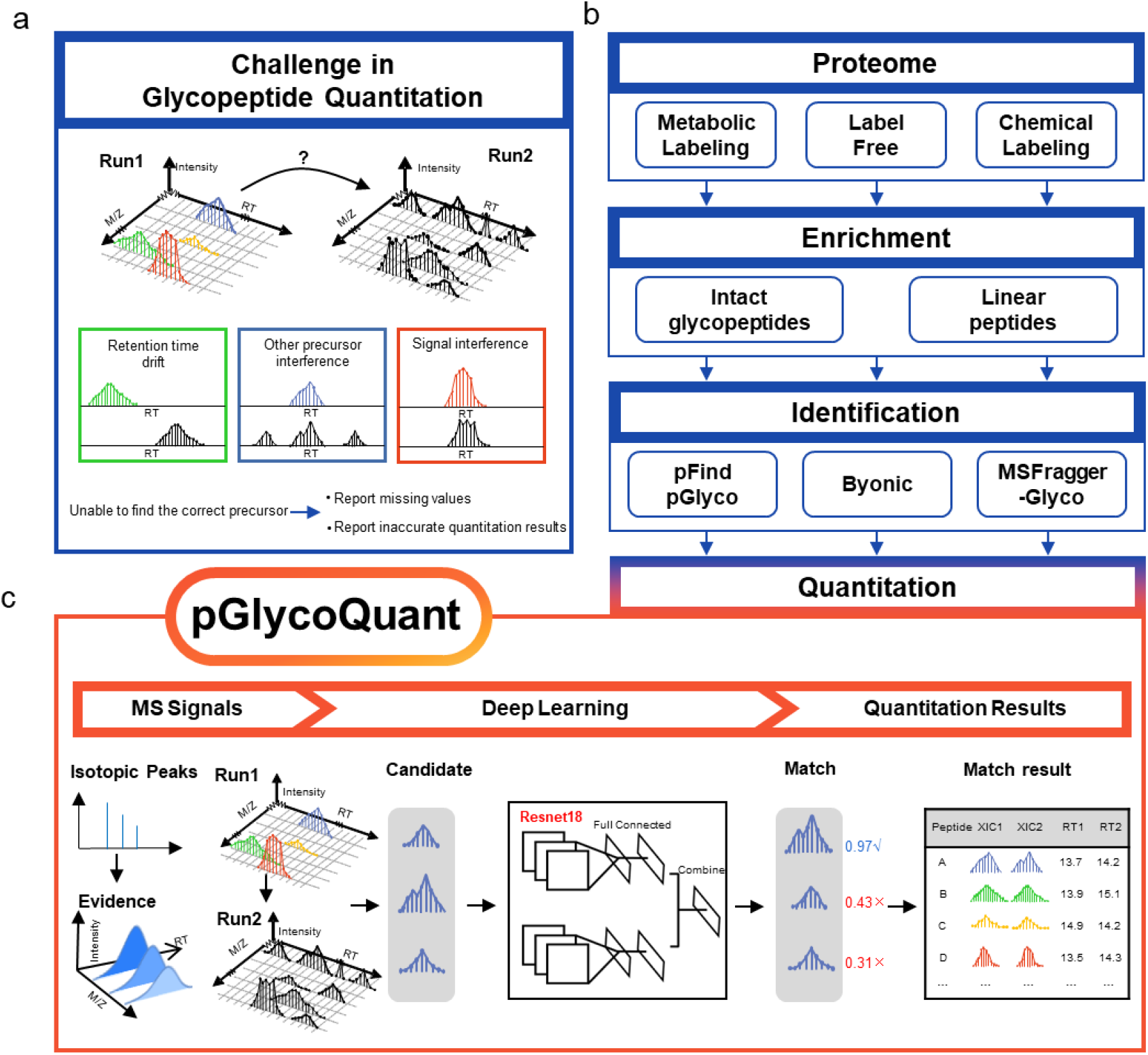
The development of pGlycoQuant. (a) Current glycopeptide quantitation software tools suffer from suboptimal reproducibility for lower-abundance signals, resulting in a high missing data rate. (b) pGlycoQuant supports both primary and tandem mass spectrometry quantitation for multiple quantitative strategies. (c) An embedded deep learning model of ResNet in pGlycoQuant.

A deep learning model is embedded in pGlycoQuant to improve the matching performance between runs (Figure 1c, Supplementary Figure 2, Online Methods). pGlycoQuant applies the ResNet deep learning to processing glycopeptide quantitative evidence between or within MS runs (Figure 1c, Supplementary Figure 2). A glycopeptide precursor is first mapped to a tensor of 1×128, and the best signal patterns are retained by the ResNet18 model. Then, a fully connected network that comprehensively utilizes multiple characteristics, including the similarity of isotopic peaks and distance of retention time between glycopeptides, is trained to measure the similarity scoring and accurate matching of two glycopeptide precursors in the same run for metabolic labeling data or between different runs for label-free data. This model also provides the matching score for each quantitation result as a softmax loss function is used in the network (Figure 1c, Supplementary Figure 2). This individual matching score evaluates the accuracy of each quantitation result (Supplementary Figure 3).

### 2. Comparative evaluation of pGlycoQuant

To evaluate the performance of pGlycoQuant for intact glycopeptide quantitation, we comprehensively compared pGlycoQuant with prevalently used search engines that equipped with quantitative functions, namely Byonic-Byologic and MSFragger-Glyco, for intact glycopeptide quantitation on three benchmark datasets, including SILAC-labeled 293T cell data, label-free HeLa cell data, and TMT-labeled mouse liver data (Figure 2a, Online Methods, Supplementary Table 1, Supplementary Table 2).

**Figure 2.**
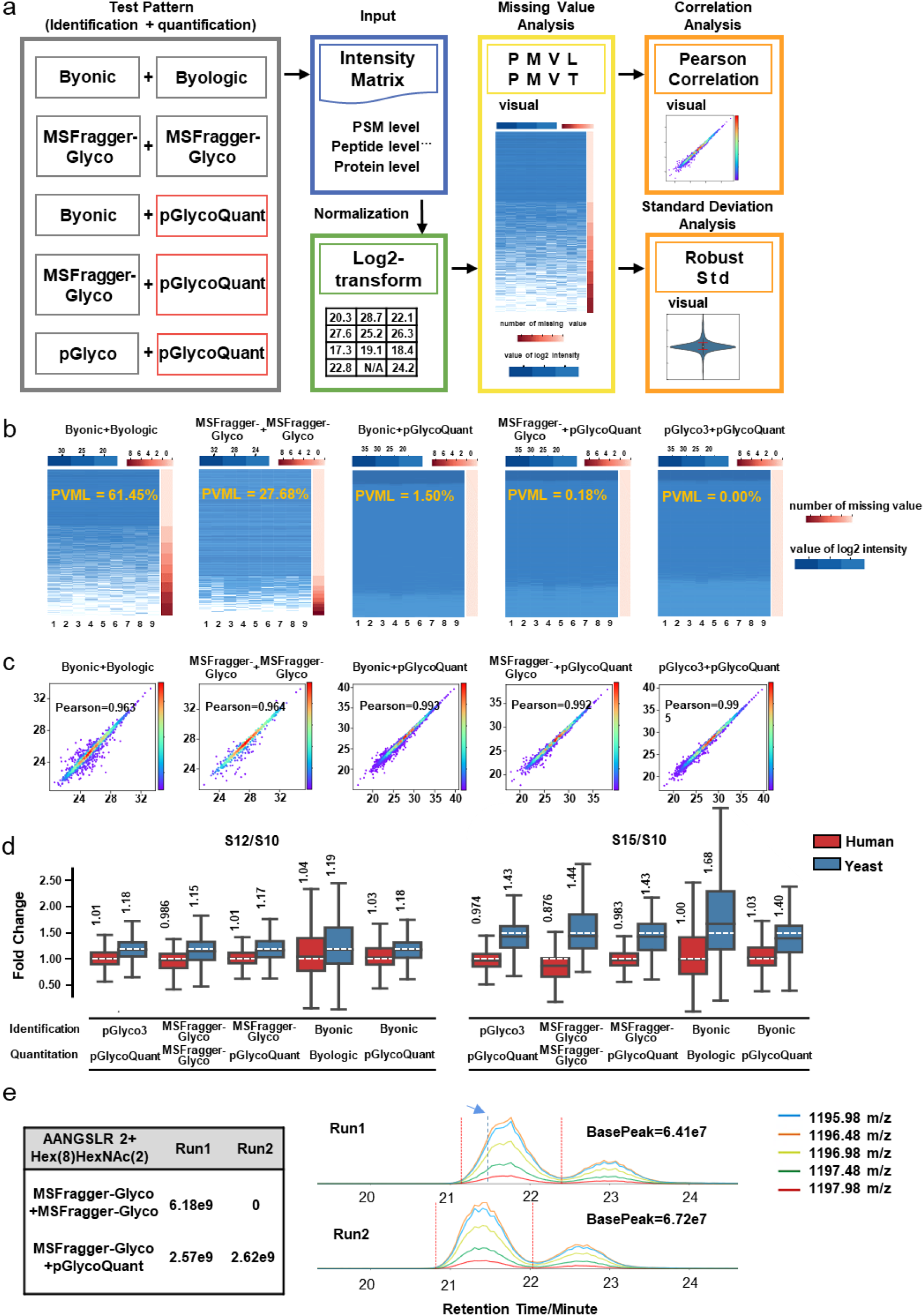
Evaluation of pGlycoQuant performance on intact glycopeptide quantitation of label-free data. (a) The comparison workflow. (b) pGlycoQuant reports few missing values, even reading the same identification results. (c) Pearson correlation of two replicate runs from the same sample. (d) Box plot visualization of the fold change of the glycopeptide quantification results of the mixed-organism samples. Percent changes were calculated based on the mean quantities in three replicates of each sample, including results with partially missing values. The medians are indicated. The boxes indicate the interquartile ranges (IQRs), and the whiskers indicate 1.5 × IQR values; no outliers are shown. The white dotted lines indicate the theoretical fold changes of the organisms (1:1:1 (S10:S12:S15) for humans and 1:1.2:1.5 (S10:S12:S15) for yeast). (e) An example of a glycopeptide signal. MSFragger-Glyco reported missing values, but pGlycoQuant correctly identified the glycopeptide signal. The table on the left shows the strength of the glycopeptide reported by MSFragger-Glyco and pGlycoQuant. The plot on the right side shows the extracted ion currents (XICs) of the glycopeptide in the mass spectrometry data, the blue dotted line represents the retention time to identify the corresponding glycopeptide by MSFragger-Glyco, and the red dotted line represents the start and end times for pGlycoQuant quantification of the glycopeptide.

Glycoproteome quantitative analysis has been hampered with the lack of high reproducibility and consistency, which is often manifested as data missing values. We defined two indicators, the proportion of missing values in line (PMVL) and the proportion of missing values in total (PMVT), to measure the proportion of missing values (Supplementary Figure 4). pGlycoQuant showed excellent coverage and extremely low proportion of missing values, while the performance of other software tools in this aspect was unsatisfactory (Supplementary Table 3). In label-free quantitation, 61.45% PMVL and 30.12% PMVT were reported by Byologic (Figure 2b, Supplementary Table 3). The missing value problem was ameliorated in MSFragger-Glyco (Figure 2b) but 27.68% PMVL and 13.45% PMVT were still reported (Supplementary Table 3). pGlycoQuant reported minuscule missing values even when reading the same GPSMs (glycopeptide spectra matches) reported by the other software tools (Supplementary Table 3). The extremely low levels of missing values obtained by pGlycoQuant can be attributed to the unique evidence generation and alignment methods (Supplementary Figure 5), which enable highly reproducible quantitation in large sample cohorts. For the SILAC-labeling data, pGlycoQuant also showed outstanding performance compared with the other tools (Supplementary Table 3). For example, for the GPSM shown in Supplementary Figure 6, pGlycoQuant reported the quantitation results according to the fixed and truly existing signals, which were ignored by other software tools. Compared to SILAC-labeling and label-free data, it is relatively easy to obtain quantitation results from TMT-labeling data, as only report ions should be considered. Unfortunately, Byologic currently does not support the quantitation of TMT-labeling data. MSFragger-Glyco could analyze the TMT-labeling data, but produced mediocre results on the PMVL and PMVT (3.77% and 1.89%, respectively, Supplementary Table 3). pGlycoQuant also reported a few missing values in TMT-labeling data analysis, which were reasonable according to the manual check (Supplementary Figure 7).

After removal of the missing values, the quantitation precision was evaluated. Because of the high PMVL and PMVT, the number of glycopeptides with quantitation values reported by other software tools was much smaller than that of pGlycoQuant after reading the same identification results. Specifically, for the hardly quantifiable glycopeptides of other software tools, pGlycoQuant provides high-precision results. pGlycoQuant could recall ∼98% of all glycopeptides without missing values (Supplementary Table 3).

Here, we use Pearson correlation and standard deviation as the measurements of precision. For SILAC-labeling data quantitation, the standard deviation of the Byologic results was almost twice as larger as that of the pGlycoQuant counterparts (Supplementary Figure 8). This is attributed to the outlier results of the low intensity glycopeptides that result in poor reproducibility of proteins with lower abundance. For the label-free and TMT-labeling data, the pGlycoQuant results reported average Pearson correlation of ∼0.99 and ∼0.91, and average standard deviation of ∼0.26 and ∼0.17, respectively (Figure 2c, Supplementary Figure 9 and Supplementary Figure 10). Other software tools also achieved comparable Pearson correlation and standard deviation, however, due to the removal of missing values during the analysis. Then, we used mixed-organism samples (human serum and budding yeast) containing two different proportions of the species^32^, to assess the precision of quantification on the basis of how well known ratios were recovered by the software tools. In comparison to Byonic and MSFragger-Glyco, pGlycoQuant demonstrated better quantification precision for both human and yeast glycopeptides (Figure 2d). By visualizing extracted ion currents (XICs) of the glycopeptide in two repeated runs of label-free data, we showed that the quantitative algorithm of pGlycoQuant can accurately locate the signal of the glycopeptide (Figure 2e).

The above results demonstrate the outstanding quantitative performance of pGlycoQuant especially on results that are difficult to quantify with software tools, thanks largely to the deep-learning-based evidence matching approaches in pGlycoQuant. Moreover, the quality control in pGlycoQuant effectively removes low-quality quantitative data, further ensuring quantitative accuracy and precision.

### 3. pGlycoQuant enabled large-scale quantitative analysis of the proteome and N-glycoproteome in different metastatic HCC cell lines

The high accuracy and precision of pGlycoQuant enable further functional exploration of site-specific glycosylation. Quantitative analyses of the proteome and intact N-glycopeptides in three HCC cell lines with different metastatic potentials (Hep3B with no metastatic potential, MHCC97L with low metastatic potential and MHCCLM3 with high metastatic potential) were performed with four replicates for each MS quantitative analysis (Figure 3a, Online Methods). A total of 11312 proteins and 11001 intact N-glycopeptides were quantified (Supplementary Data 1, 2), among which those that appeared more than in duplicate were regarded as reliable. The results showed that a total of 9154 proteins and 6435 intact N-glycopeptides were reliably identified and quantified from the proteomic and intact N-glycopeptide quantitation experiments, respectively (Figure 3b, Supplementary Data 1, 2), which was the largest intact glycopeptide quantitative result in the three cell lines thus far. The 6435 intact N-glycopeptides were attributed to 769 glycoproteins with 1357 N-glycosites and 143 N-glycans (Figure 3c). The quantitative ratios among the three cell lines showed high correlation, demonstrating reliable quantitative accuracy and good repeatability (Supplementary Figure 11,12). The criteria of ratio ≥2 or ≤0.5 and p<0.01 were adopted as significant differential expression to further filter the quantitative results, resulting in 2438 proteins and 3030 intact glycopeptides. The details of the differential proteins and intact glycopeptides among cell lines are listed in Supplementary Data 3, 4.

**Figure 3.**
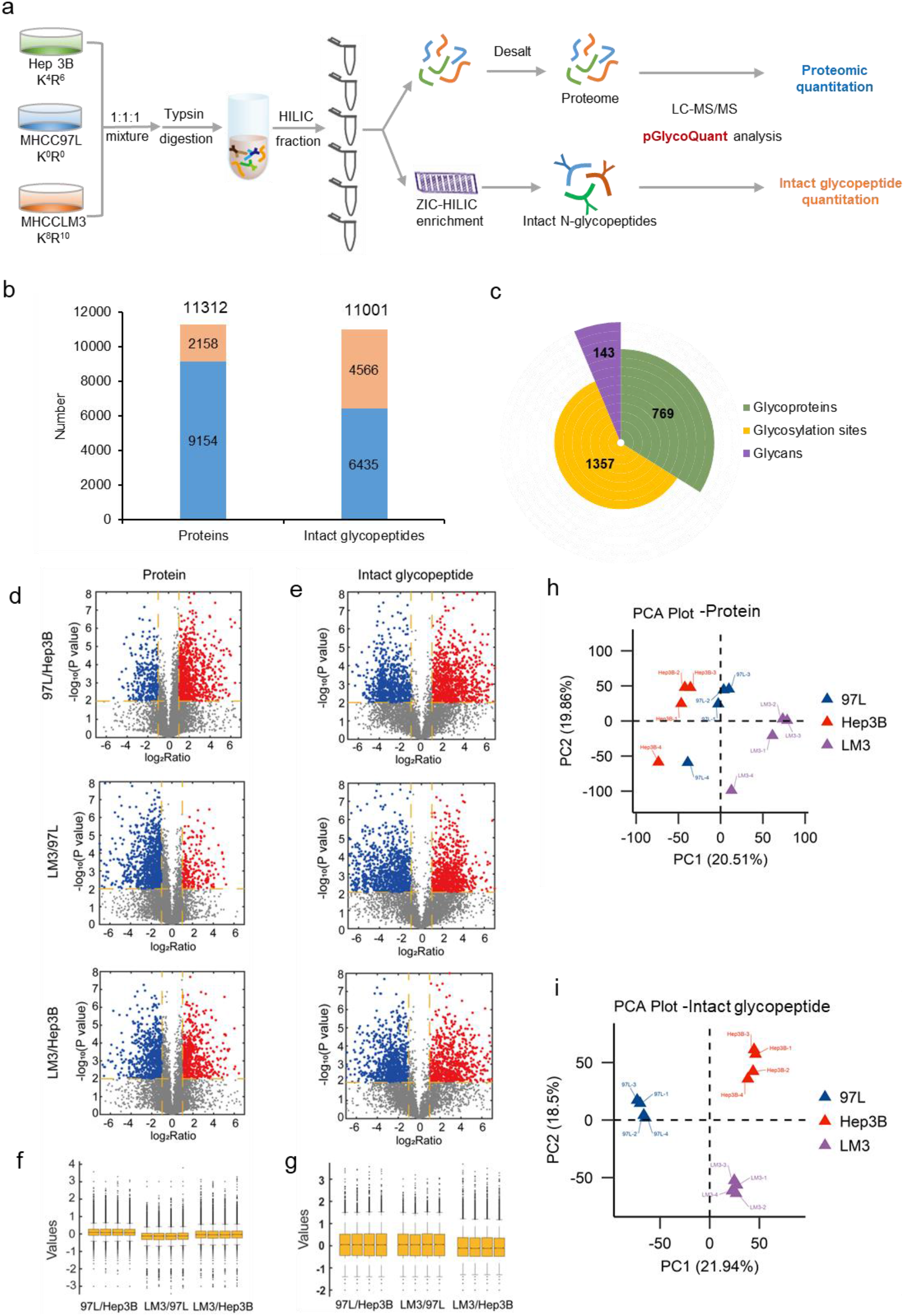
Large-scale quantitative analyses of the proteome and N-glycoproteome of different metastatic HCC cell lines with pGlycoQuant. (a) Experimental workflow. (b) The numbers of proteins and intact glycopeptides quantified from proteomic and intact N-glycopeptide quantitation experiments, respectively (the blue bar represents the number of proteins/glycopeptides quantified in more than two replicates). (c) General information on 6435 reliably quantified intact N-glycopeptides. Volcano map of differential proteins (d) and intact glycopeptides (e) among three cell lines. Box diagram with four replicates for the quantitation of proteins (f) and intact glycopeptides (g). Principal component analysis (PCA) plot of differential proteins (h) and differential intact glycopeptides (i) among three cell lines.

The ability to quantify the proteome and intact glycopeptides at such a large scale provides opportunities to investigate the role of glycosylation in the metastasis of HCC. Gene Ontology (GO) analyses of the proteome and glycoproteome showed that differential proteins were mainly concentrated in the cytoplasm and nucleus (Supplementary Figure 13a), were associated with ion binding and RNA/DNA binding (Supplementary Figure 13b), and participated in cellular metabolism and signal transduction (Supplementary Figure 13c), while differential intact N-glycopeptide-related glycoproteins were more likely to be located in the membrane and extracellular regions (Supplementary Figure 14a), to be related to enzyme binding and hydrolase activity (Supplementary Figure 14b), and to be involved in cell adhesion(Supplementary Figure 14c).

Then, we used volcano plots and box plots to visually show the distribution and dispersion degree of the differential proteins and intact glycopeptides. The differences in glycopeptides were more diffuse than those in proteins (Figure 3d-g) in the three cell lines. Further principal component analysis (PCA) showed that compared to proteomes, differences in intact glycopeptides were more likely to distinguish the three HCC cell lines with different metastatic potentials (Figure 3h, i). Thus, we drilled down for in-depth information on intact glycopeptide data to explore the role of site-specific N-glycosylation in the metastasis of HCC.

### 4. Site-specific N-glycoproteomic analyses revealed great heterogeneity and implied altered core fucosylation to be highly associated with in vitro cell invasion and metastasis

Statistical analyses of the site-specific N-glycoproteome enable further visualization of glycoproteome heterogeneity and investigation of system-wide glycosylation patterns^33^. Firstly, we performed overall statistical analyses on site-specific N-glycoproteome data from three cell lines. It was demonstrated that ∼79.5% of the glycosites (1079 of the 1357) quantified in this study were annotated in the UniProt database (Figure 4a). In addition to quantifying 456 previously proven glycosties (456 published and 48 imported), we provided experimental evidence for 575 UniProt-predicted glycosites (572 sequence analyses and 3 by similarity) and 278 non-UniProt-recorded glycosites (Figure 4a). Sequence motif analysis showed that the majority of N-glycosites share N-X-S (40%) and N-X-T (58%) sequons, while only 2% of the glycosites have the N-X-C sequon (Figure 4b). The distribution of singly or multiply glycosylated proteins and the degree of glycan microheterogeneity showed that more than half of the glycoproteins (481 of the 769 identified glycoproteins) had only one glycosite (Figure 4c), while 75% of glycosites contained more than one glycan (Figure 4d). Glycans with 8-12 monosaccharides dominated in these data (Figure 4e). A network between glycan types and glycosites on glycoproteins revealed that the fucosylation type was prevalent in the HCC cells, and fucosylation and sialylation occurred more frequently on multiply-glycosylated proteins, thus contributing more to heterogeneity (Figure 4f). A heatmap displaying the frequency of glycan pairs co-occurring at the same site illustrated that oligomannose appears to co-occur with several groups of complex/hybrid, fucosylation and sialylation types with high frequency (Figure 4g), which further indicates site-specific microheterogeneity.

**Figure 4.**
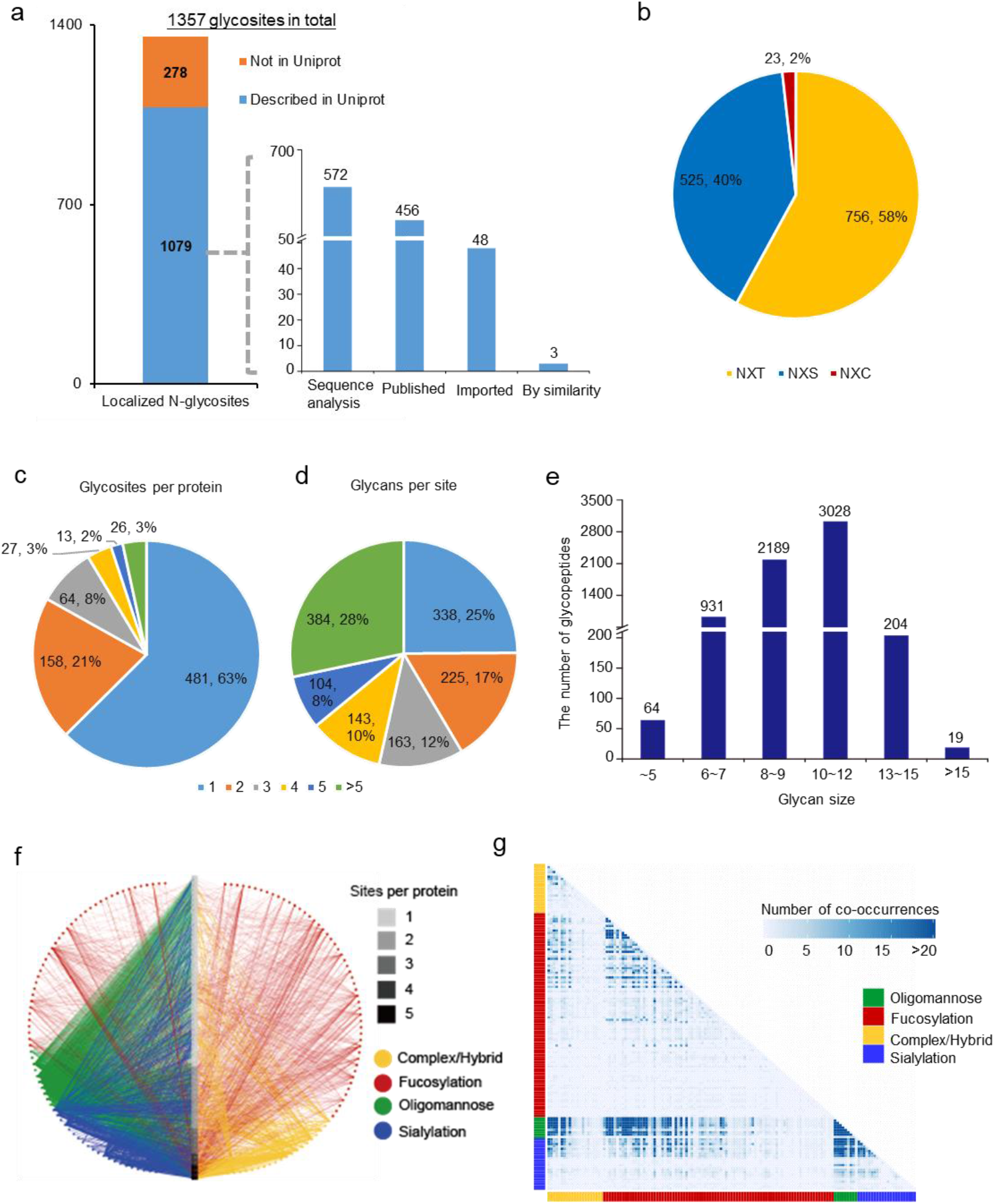
Characteristics of site-specific N-glycans quantified in HCC cell lines. (a) The status of quantified glycosites recorded in the UniProt database. (b) Recognition of the sequence motif of N-glycosylation (N-X-S/T/C, where N is asparagine, X is any amino acid except proline, S is serine, T is threonine, and C is cysteine.) (c) The number and percentage of N-glycosites located on a certain protein. (d) The number and percentage of N-glycans linker to a certain glycosite. (e) The glycan size distribution in 6435 glycopeptides from three HCC cell lines. (f) Glycoprotein-glycan network maps of specific glycans (outer circle, 143 in total) modifying specific glycoproteins (inner bar, 769 in total). (g) A glycan co-occurrence heatmap representing the number of times glycan pairs appear together at the same glycosite, indicating which glycans contribute most to the microheterogeneity of the 1019 glycosites with more than one glycan modifying them.

We further analyzed different distributions of glycan size and glycan type in all quantified intact glycopeptides and the uniformly up-/downregulated glycopeptides in three cell lines with increased metastatic potential. It could be concluded that upregulated glycopeptides tend to have longer glycans (mostly with 8-12 monosaccharides) than downregulated glycopeptides (Supplementary Figure 15a). Comparing glycan types in all quantified glycopeptides showed that fucosylation and sialylation were more associated with upregulated glycopeptides, while oligomannose was dominant in downregulated glycoppetides (Supplementary Figure 15b).

Glycosyltransferases (GTs) and glycoside hydrolases (GHs) coregulate the synthesis of glycans and are key factors affecting protein glycosylation. Thus, we then analyzed glycan-related enzymes. We quantify 164 glycan-related genes, including 85 GTs and 44 GHs, from the proteome quantitative results of 9154 proteins (Supplementary Figure 16, Supplementary Data 5). Among them, we noted that a GT that regulates core fucosylation synthesis, alpha-(1,6)-fucosyltransferase (FUT8), was significantly changed in the three cell lines (Supplementary Figure 16), which implies that core fucosylation is highly correlated with HCC cell metastasis.

### 5. Site-979-specific core fucosylation of L1CAM was identified and validated in vitro as a potential regulator of HCC metastasis

Consequently, we further analyzed core-fucosylated glycoproteins, and our screen identified a glycoprotein, L1CAM, in which glycosite 979 is highly core-fucosylated and upregulated in three cell lines with increasing metastatic potential. A total of 35 site-specific glycans, including 5 glycosites and 20 glycans were quantified in L1CAM (Figure 5a, b). L1CAM is a highly glycosylated protein known to regulate cell attachment, invasion and migration in several cancers and is associated with poor prognosis^34-36^. For example, Mahal and Hernando et al.^37^ demonstrated that glycoprotein targets of FUT8 were enriched in cell migration proteins, including the adhesion molecule L1CAM, in melanoma metastases. However, little is known about the site-specific glycosylation of L1CAM associated with cell invasion and metastasis. We found that core fucosylation with glycan composition Hex[5]HexNAc[4]NeuAc[1]Fuc[1] at glycosite 979 was significantly high in L1CAM in all three cell lines (Supplementary Figure 17a) through normalization within one cell line. After normalization among the three cell lines, it was obvious that all fucosylated glycans at site 979 of L1CAM were consistently upregulated with increasing metastatic potential of the cell lines (Supplementary Figure 17b). Further analysis of the protein expression levels of L1CAM and FUT8 from proteomic quantitation data revealed that the upregulated of fucosylation at site 979 of L1CAM with increasing metastatic potential was caused by different reasons (Figure 5c): from no metastatic potential to low metastatic potential, the increased fucosylation at site 979 was caused by the increased expression of FUT8; from low metastatic potential to high metastatic potential, the increased fucosylation at site 979 was mainly due to the increased protein content of L1CAM. The western blot results were consistent with the of MS-based proteomic quantitative results interpreted by pGlycoQuant (Figure 5d-f). Based on the above results and the known ability of L1CAM support invasion and metastasis, we hypothesize that increased core fucosylation at site 979 of L1CAM reduces L1CAM cleavage by plasmin, facilitating HCC cell line invasion and metastasis (Figure 5g).

**Figure 5.**
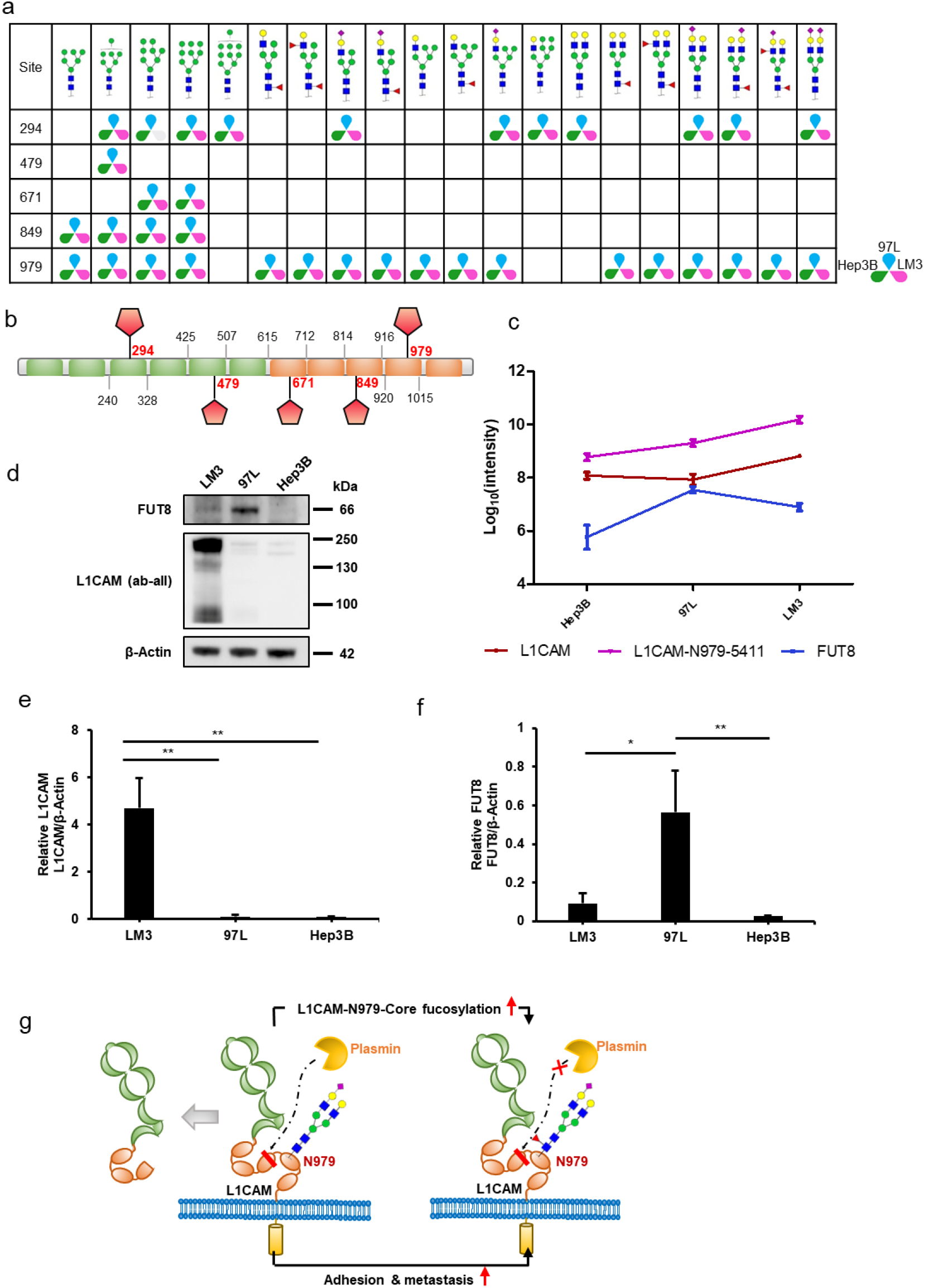
Site-979-specific core fucosylation of L1CAM is upregulated in three HCC cell lines with increasing metastatic potential. (a) Site-specific N-glycosylation of L1CAM in three HCC cell lines. (b) Graphical view of five identified glycosites located in specific positions and domains of L1CAM. The orange bar chart represents the Ig-like C2 type, and the green bar chart represents fibronectin type III. (c) Change trends of protein L1CAM, site-979-specific glycan Hex[5]HexNAc[4]NeuAc[1]Fuc[1], and protein FUT8 in three HCC cell lines from low metastatic potential to high metastatic potential. (d) Western blot validation of the protein expression levels of L1CAM and FUT8 in three HCC cell lines. (e) L1CAM levels in three HCC cell lines. (f) FUT8 levels in three HCC cell lines. (g) Hypothesis of increased core fucosylation at site 979 of L1CAM facilitating HCC cell line invasion and metastasis. The glycan symbols are as follows: green circle for Hex, blue square for HexNAc, purple diamond for sialic acids and red triangle for fucose.

We performed several experiments to investigate the impact of site-979-specific core fucosylation of L1CAM on in vitro HCC cell metastasis. We first examined whether L1CAM is required for the maintenance of existing metastasis by the silencing of L1CAM in LM3 cells. Consistently, siL1CAM cells displayed decreased L1CAM protein (Supplementary Figure 18a) and reduced cell migration and invasion in comparison to siCtrl cells (Supplementary Figure 18b-d). To confirm the role of site-979-specific core fucosylation of L1CAM in in vitro HCC cell metastasis, we next investigated whether L1CAM overexpression with or without the site-979 mutation has the same ability to promote HCC cell metastatic capacity and whether the core fucosylation of L1CAM contributes to that effect. L1CAM overexpression triggered significant increases in 97L cell migration and invasion in vitro, while site-979-mutated L1CAM overexpression with the same protein amount showed no prometastatic effects (Figure 6a-d). Further silencing of FUT8 in 97L cells (Figure 6e), which resulted in reduced core-fucosylated L1CAM (Figure 6f), decreased in vitro cell migration and invasion (Figure 6g-i;). These results suggest that site-979-specific core fucosylation is critical to prometastatic phenotype in HCC cell lines. Previous studies have reported that the cleavage of L1CAM by plasmin inhibits its ability to mediate neural cell invasion and metastatic outgrowth^38, 39^. Here, we observed that 97L cells with site-979-mutated L1CAM overexpression indeed tended to be more easily cleaved by plasmin than unmutated cells (Figure 6j), which to a certain extent accounted for the impact of altered core fucosylation at site 979 of L1CAM on L1CAM cleavage by plasmin.

**Figure 6.**
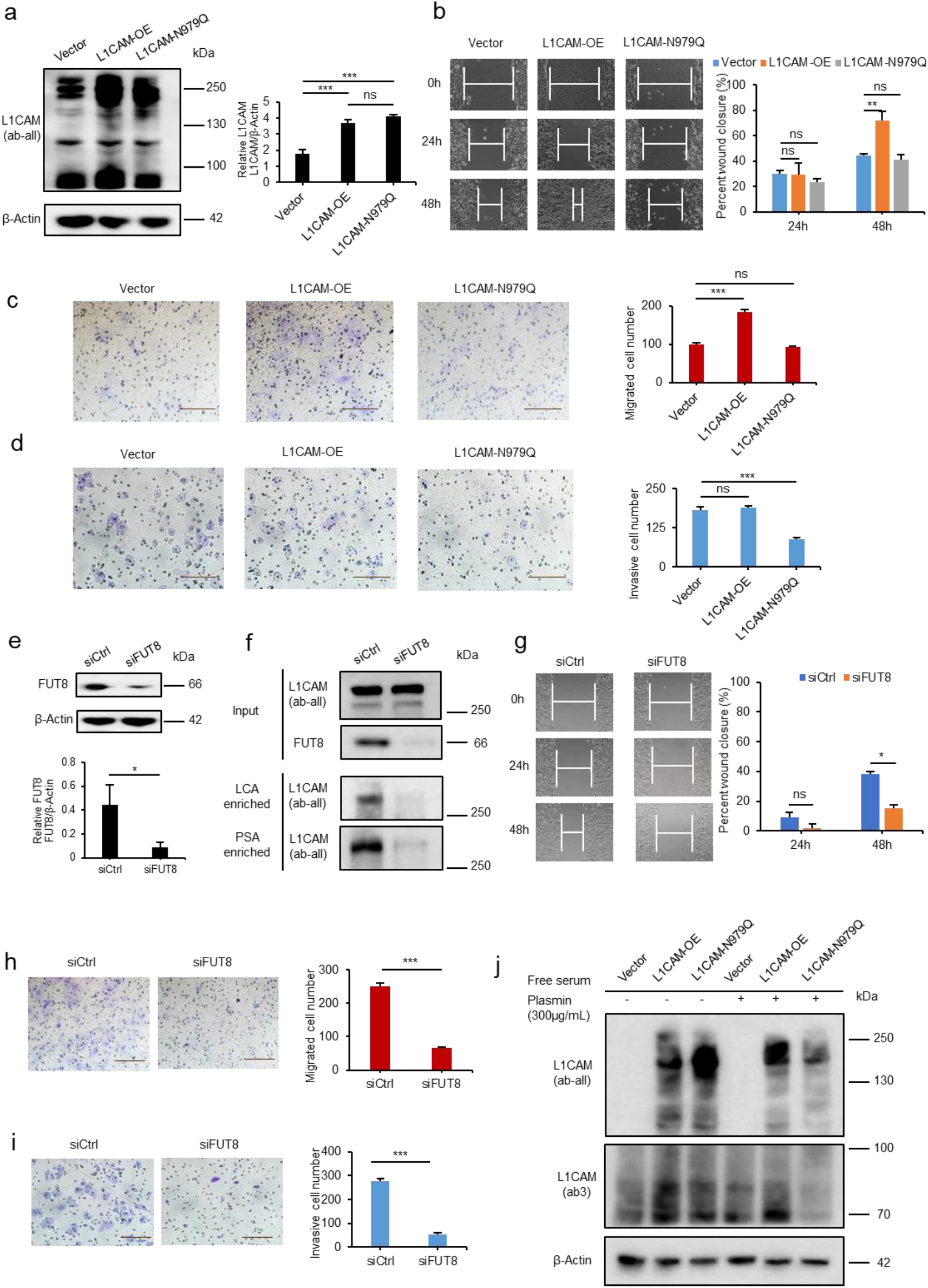
In vitro validation of site-979-specific core fucosylation of L1CAM is a potential regulator of HCC metastasis. (a) Western blot of L1CAM levels in 97L cells overexpressing control vector, pL1CAM-FLAG vector or pL1CAM (N979Q)-FLAG vector. (b) Wound healing assay in monolayers of 97L-overexpressing cells (vector, L1CAM-OE and L1CAM-N979Q). The scratch area of the cells was detected with an inverted microscope (10X). (c) Transwell migration assay and (d) Matrigel invasion assay (scale bar = 100 µm) in 97L-overexpressing cells (vector, L1CAM-OE and L1CAM-N979Q). (e) Western blot of L1CAM levels in 97L cells transfected with negative control (Ctrl) and FUT8 siRNA. (f) Lectin enrichment in the lysate of 97L cells transfected with Ctrl or FUT8 siRNA followed by western blot with anti-L1CAM (ab-all) antibody. Input showed no effect of FUT8 knockdown on L1CAM expression. (g) Wound healing assay in monolayers of 97L-knockdown cells (siCtrl and siFUT8). The scratch area of the cells was detected with an inverted microscope (10X). (h) Transwell migration assay and (i) Matrigel invasion assay (scale bar = 100 µm) in 97L-knocknown cells (siCtrl and siFUT8). (j) Western blot of cleaved L1CAM (ab3) and full-length L1CAM (ab-all) in whole lysates of 97L-overexpressing cells (vector, L1CAM-OE and L1CAM-N979Q) after treatment with plasmin. Cells were transfected for 24 h and then incubated with plasmin (300 μg/mL) for 8 h. The grayscale values of the western blot data (a, e) and scratch area of the wound healing assay were measured by Image J. β-Actin was used for the normalization of loading in all western blot data (a, e). The data shown are representative of three independent experiments and are presented as the means ± SD. P values were determined by the two-tailed unpaired t-test. ∗p < 0.05, ∗∗p < 0.01, and ∗∗∗p < 0.001 compared to the control (n = 3).

## Discussion

Since the identification of intact glycopeptide has been greatly facilitated by software tools^5, 40, 41^, there is an urgent need to develop efficient tools for accurate intact glycopeptide quantitation to assist in exploring differences in site-specific glycosylation^6, 12^. The main challenge in accurate quantitation by LC–MS-based methods is to correctly extract the targeted mass spectral signal since it is easily interfered with nearby signals or noise. Herein, we developed pGlycoQuant to support multiple common glycopeptide quantitative strategies. pGlycoQuant applies a ResNet deep learning model to process glycopeptide quantitative evidence between or within MS runs. The ResNet model can learn the in-depth representation of glycopeptide quantitative evidence in complex mass spectrometry data, improving the sensitivity and precision in detecting low-abundance glycopeptides signals. Moreover, the deep-learning model reports the matching score, which is difficult to measure by the traditional algorithms. We benchmarked our pGlycoQuant with several prevalently used software tools on three different quantitation strategy-based datasets, and the results demonstrated that pGlycoQuant outperforms other tools in terms of precision and reproducibility.

Precise and minuscule-value glycoproteome quantitation with pGlycoQuant at the site-specific glycosylation level provides us with new opportunities and horizons to explore the role of glycosylation organisms. The combination of large-scale quantitative analyses of the proteome and glycoproteome in three different metastatic HCC cell lines demonstrates a generic application of pGlycoQuant for investigating the role of site-specific glycosylation, yielding the largest intact glycopeptide quantitative data in three HCC cell lines and enabling the visualization of glycoproteome heterogeneity and the investigation of system-wide glycosylation patterns. Based on the convincing quantitative results obtained by pGlycoQuant, fortunately, the site-979-specific core fucosylation of L1CAM was identified in a screen and validated as a potential regulator of HCC metastasis in vitro, which presents the necessity and possibility of pGlycoQuant in biological research.

Currently, pGlycoQuant is compatible with many search engines, including pFind, pGlyco2.0, pGlyco3, MSFragger-Glyco, and Byonic, providing a convenient way to quantify the glycoproteome at the site-specific level for the majority of users. Although pGlycoQuant is shown here in the context of N-glycoproteomic quantitation, it is also applicable to intact O-glycopeptide quantitation. With a deep residual network for precise reporting with minuscule missing values, pGlycoQuant makes it possible to quantitatively investigate site-specific glycosylation and illuminate its functions.

## Online Methods

### Workflow of pGlycoQuant

As shown in Supplementary Figure 1, the workflow of pGlycoQuant consists of three steps:

#### Step 1

Reading the identification results. pGlycoQuant can read the identification results from pGlyco, Byonic and MSFragger-Glyco. High confidence peptide-spectrum matches (PSMs) produced by identification software tools are read into the program.

#### Step 2

Extracting the quantitation signals. For each input PSM, pGlycoQuant calculates the theoretical distribution of isotopic peaks using a stepwise convolution algorithm and identifies experimental isotopic peaks in a range of MS scans where the peptide may be expected. pGlycoQuant constructs chromatograms for individual isotopic peaks of the peptides^17^. These “isotopic chromatograms” are called the “feature” in other papers, but this word is confused with the computational word “feature” in machine learning, so they are called the “evidence” of a peptide in this paper. For the chemical labeling data, pGlycoQuant picks the reported ion peaks in MS2 scans according to the input parameters.

#### Step 3

Quantitation of peptide and protein intensities. (1) Metabolic label data. The chromatogram area of light and heavy peptides is recorded as the peptide intensity, and the sum of all corresponding peptide intensities is the protein intensity. Furthermore, pGlycoQuant calculates the similarity score based on the well-trained deep learning model (see the additional step below) to measure the accuracy of the quantitation result. (2) Label-free data. Unlike metabolic label quantitation, for each identified peptide in one run, pGlycoQuant detects the corresponding evidence in other runs, i.e., matches evidence between runs. Given an identified peptide in one run, pGlycoQuant constructs its evidence in the full MS scans and then calculates the similarity scores of all the evidence with the same precursor mass in a ±2 min. (user defined) retention time window in another run. This similarity score of two pieces of evidence is calculated based on the same well-trained neutral network in metabolic label quantitation, but the pair of pieces of evidence with the maximum similarity score is selected to obtain the quantitation result between different runs. (3) Chemical label data. The peptide and protein intensities are calculated by summing the intensities of the reported ion peaks of the corresponding PSMs.

#### Additional Step

Training a deep-learning-based evidence matching model. As illustrated in Supplementary Figure 2, we use label-free data to train the deep learning model. A total number of 3000 high score peptides simultaneously identified in two runs with very similar retention times were selected, and the pairs of evidence are defined as positive samples. A total of 3000 peptides identified in only one run are selected, and the evidence in the identified run and random evidence in another run with different precursor masses are defined as negative samples. These positive and negative samples are used to train the following deep learning model.

We consider the evidence as a matrix, similar to a picture in computational vision. The matrix is then transformed to a 512*1 vector by the ResNet18 model. This transformation is the critical operation to measure the pattern of a peptide evidence. The two vectors from two pieces of evidence are then combined into a 1024*1 vector, which is the input of a fully connected neural network. This network is designed to describe the similarity of the two original pieces of evidence, and a 16*1 vector is its output. Moreover, given a pair of pieces of evidence, 10 classical features are also extracted as a 10*1 vector. Another fully connected neural network with a softmax loss function is designed to output the final matching score of the two original pieces of evidence. This final matching score, called the similarity score in this paper, is in the interval of [0, 1], where 0 corresponds to very dissimilar and 1 to very similar evidence.

### Comparison of pGlycoQuant with other N-glycoproteome quantitation software tools

To guarantee a fair comparison, we adopted the following procedure based on previously suggested rules: (1) To prevent differences introduced by identification, pGlycoQuant reads the identification results reported by other software tools and calculates the quantitation results. (2) Missing values are analyzed, and two proportions at the protein level and protein quantitation value level are reported. (3) After the removal of the missing values, we compare the Pearson correlation and standard deviation of the intensities at the PSM, glycopeptide and protein levels without normalization. The key indicators are listed below (see Supplementary Figure 4):

PMVL (proportion of missing value in line) = No. peptides with more than one missing value/No. all peptides

PMVT (proportion of missing value in total) = No. individual missing values/No. all quantitation values

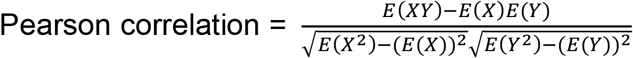, where X and Y are vectors of protein or peptide intensities in one run, respectively.

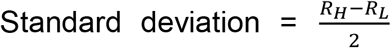, where R is the vector of log2-transformed quantitation ratios of two runs. R_H_ and R_L_ are the 84.13% and 15.87% percentiles, respectively. This robust standard deviation was introduced in the MaxQuant paper. For a normal distribution, these would be equal to each other and to the conventional definition of a standard deviation.

### Quantitative and comparative analyses of intact N-glycopeptide results from three benchmark datasets

Three benchmark datasets, namely, SILAC-labeled 293T cell data, label-free HeLa cell data, and TMT-labeled mouse liver data, were generated with the different quantitative strategies. In brief, for the SILAC-labeled 293T data, 293T cells were cultured in K0R0 and K6R6 media. Then, proteins were extracted from the labeled cells, mixed at a 1:1 ratio and digested. For the label-free HeLa data, HeLa cells were directly collected and used for protein extraction and digestion. For the TMT-labeled mouse liver data, proteins were extracted from ground mouse livers and digested. The digests were divided into two aliquots, each of which was labeled with the TMT6plex™ label reagents TMT^6^-127 and TMT^6^-130, respectively, following the TMT6plexTM isobaric label reagent product manual (Thermo Fisher Scientific, Waltham, MA, U.S.A.), and mixed with 1:1. Finally, as with any quantitative strategies, glycopeptides were enriched from the desalted digests using ZIC-HILIC method and analyzed by LC–MS/MS. Detailed sample preparation and data acquisition methods are described in Supplementary Note 1.

### Quantitative analyses of the proteome and N-glycoproteome in three HCC cell lines with SILAC labeling

We used the SILAC strategy to label the three cell lines MHCC97L, Hep3B and MHCCLM3 with K0R0, K4R6 and K8R10 labeling, respectively. Then, proteins were extracted from the labeled cells, mixed at a 1:1:1 ratio and digested. The tryptic digests were then subjected to chromatographic fractionation with HILIC and used for direct proteomic quantitation and intact N-glycopeptide quantitation after ZIC-HILIC enrichment by four replicates of LC–MS/MS analysis. See Supplementary Note 1 for details. For the SILAC labeling, cells were cultured following the experimental procedure described in Supplementary Note 1 and collected after culturing for 8 generations with over 95% labeling efficiency. To confirm the performance of our SILAC labeling experiments, we mixed the proteins from different labeling cells at a 1:1:1 ratio, digested them and quantitatively analyzed them by LC– MS/MS. We used housekeeping proteins, including actin, tubulin and GAPDH, which are usually stable in organisms, as standards to evaluate the labeling efficiency. All the relative quantitative results of these housekeeping proteins showed no significant changes among the three cell lines (Supplementary Data 3), which demonstrated a good SILAC experiment and guaranteed the feasibility of further quantitative analysis.

### Benchmark, software versions

pGlycoQuant supports quantitation of the identification results from Byonic, MSfragger, pFind and pGlyco services. The following search engines and quantitation engines/modes were used in this study for the pGlycoQuant-supporting quantitation test and comparison. Search engines: pGlyco3, MSFragger-Glyco, and Byonic. Quantitation engines/mode: pGlycoQuant, MSfragger-Glyco, and Byologic. The detailed versions are listed in Supplementary Table 1.

### Database searching

Three benchmark datasets, including SILAC-labeled 293T cell data, label-free HeLa cell data, and TMT-labeled mouse liver data, were searched using different software tools (Supplementary Table 1) for quantitative performance comparison. The detailed searching parameters are shown in Supplementary Table 2. The proteome data of SILIAC-labeled HCC cell lines were analyzed using Open-pFind software^28^ with open search mode for identification followed by pGlycoQuant for quantitation. The intact glycopeptide data of SILIAC-labeled HCC cell lines were analyzed using pGlyco3 followed by pGlycoQuant for quantitation with the same parameters in Supplementary Table 2.

### In vitro functional validation and molecular biology experiments

We utilized western blotting to detect the expression of L1CAM and FUT8 in the three cell lines Hep3B, MHCC97L, and MHCCLM3 and verify MS-based proteomic quantitative results. The 97L cells were transfected with FUT8 siRNA, pL1CAM-FLAG plasmid and pL1CAM (N979Q)-FLAG plasmid, and the LM3 cells were knocked down by L1CAM siRNA. The effect of transfection was tested through western blot or lectin enrichment and immunoblot assays. For functional validation experiments, wound healing assays and transwell migration assays were adopted to validate the migration capacity of the above transfected cells. The invasive ability of these cells was evaluated by Matrigel invasion assay. The details of the above experiments are described in Supplementary Note 2.

## Acknowledgements

We thank Professor Simin He from Institute of Computing Technology, CAS, Beijing, China for his kindly directing research, providing valuable advices and moral support. This work was supported by grants from the National Natural Science Foundation of China Project (91853102 to W.C.), and the innovative research team of high-level local university in Shanghai. This paper is dedicated to the memory of Professor Pengyuan Yang (1949.6.12–2021.5.31), who passed away during the paper preparation.

## Data availability

The RAW MS data generated in this work, as well as the search results and relevant analyses will be exposed, once the paper is published.

## Code availability

pGlycoQuant could be downloaded from https://github.com/expellir-arma/pGlycoQuant/releases/

## Author Contributions

W.C. conducted this project, performed the wet-lab experiments and data analysis, and wrote the manuscript. C.L. developed the software pGlycoQuant, performed the data analysis and wrote the manuscript. S.K. conducted the experiments for molecular function validation and revised the manuscript. W.Z. contributed to the pGlycoQuant development and revised the manuscript. B.Y. contributed to the wet-lab experiments for proteome and glycoproteome identification in HCC cell lines. P.G. and X.H. contributed to the pGlycoQuant development and data analysis. H.Z. and G.Y. contribute to the LC-MS/MS analysis. Y.Z., M.L., X. Q., and M. W. contributed to the MS data analysis. W. C, C.L., and P.Y. supervised this project.

