## Supplementary Information for "pGlycoQuant with a deep residual network for precise and minuscule-missing-value quantitative glycoproteomics enabling the functional exploration of site-specific glycosylation"

### Supplementary Table 1 Sources of Data and software tools

#### 1-1 Sources of data

| Data | Sample Source | Raw data Type | Raw data number |
|---|---|---|---|
| SILAC-labeled 293T cell data | 293T cell | SILAC | 3 |
| label-free Hela cell data | Hela cell | label-free | 9 |
| TMT-labeled mouse liver data | mouse liver | TMT-labeled | 1 |

#### 1-2 Sources of software tools

| Quantification software | Identification software | Version of Identification software | Quantification software | Version of Quantitation software |
|---|---|---|---|---|
| Byologic | Byonic | V4.0.1 | Byologic | V4.0.1 |
| pGlycoQuant | Byonic | V4.0.1 | pGlycoQuant | 1.1 |
| MSFragger-Glyco+ MSFragger-Glyco | MSFragger-Glyco | FragPipe15.0, MSFragger_3.2, philosopher_v3.3.12 (SILAC-labeled 293T cell data and label-free Hela cell data) | MSFragger-Glyco | FragPipe15.0, MSFragger_3.2, philosopher_v3.3.12 (SILAC-labeled 293T cell data and label-free Hela cell data) |
| | | FragPipe16.0, MSFragger_3.3, philosopher_v4.0.0 (TMT-labeled mouse liver data) | | FragPipe16.0, MSFragger_3.3, philosopher_v4.0.0 (TMT-labeled mouse liver data) |
| MSFragger-Glyco+ pGlycoQuant | MSFragger-Glyco | FragPipe15.0, MSFragger_3.2, philosopher_v3.3.12 (SILAC-labeled 293T cell data and label-free Hela cell data) | pGlycoQuant | 1.1 |
| | | FragPipe16.0, MSFragger_3.3, philosopher_v4.0.0 (TMT-labeled mouse liver data) | | |
| pGlyco3+pGlycoQuant | pGlyco3 | pGlyco3 | pGlycoQuant | 1.1 |

### Supplementary Table 2 Parameters of all software tools

#### 2-1 SILAC-labeled 293T cell data

| **Dataset** | | SILAC-labeled 293T cell data | | | | |
|---|---|---|---|---|---|---|
| **Software** | Analysis Software | pGlyco3+pGlycoQuant | Byonic+Byologic | Byonic+pGlycoQuant | MSFragger-Glyco+ MSFragger-Glyco | MSFragger-Glyco+pGlycoQuant |
| | Identification Software | pGlyco3 | Byonic | | MSFragger-Glyco | |
| **Peptide** | Protein database | Human-H.sapiens-SP-1808.fasta | | | | |
| | Digestion | trypsin | trypsin | | trypsin | |
| | Max miss cleavage | 2 | 3 | | 2 | |
| | Fixed Modification | Carbamidomethyl on Cys | Carbamidomethyl on Cys | | Carbamidomethyl on Cys | |
| | Variable Modification | Acetyl on Protein N-Term/Oxidation on Met | Acetyl on Protein N-Term/Oxidation on Met | | Acetyl on Protein N-Term/Oxidation on Met | |
| **Modifications** | Mode | N-glycan | N-glycan | | N-Glycan | |
| | Glycan database | N-Human | 182 Human | | 182 Human | |
| **QC** | GPSM filter | 1%FDR | Score≥200&\|logProb\|≥2 | | 1%FDR | |
| **MS** | Fragmentation Method | HCD | HCD | | HCD | |
| | Precursor tolerance | 4ppm | 4ppm | | 20ppm | |
| | Fragment tolerance | 20ppm | 20ppm | | 20ppm | |
| **Quantification** | Quantification Software | pGlycoQuant | Byologic | pGlycoQuant | MSFragger-Glyco | pGlycoQuant |
| | Quant type | SILAC | | | | |
| | Quant level | MS1 | MS1 | MS1 | MS1 | MS1 |
| | Precursor tolerance | 20ppm | — | 20ppm | 10ppm | 20ppm |
| | Fragment tolerance | — | | | | |
| | RT tolerance window | 2min | — | 2min | — | 2min |

### 2-2 label-free Hela cell data

| **Dataset** | label-free Hela cell data | | | | | |
|---|---|---|---|---|---|---|
| **Software** | Analysis Software | pGlyco3+pGlycoQuant | Byonic+Byologic | Byonic+pGlycoQuant | MSFragger-Glyco+ MSFragger-Glyco | MSFragger-Glyco+pGlycoQuant |
| | Identification Software | pGlyco3 | Byonic | | MSFragger-Glyco | |
| **Peptide** | Protein database | Human-H.sapiens-SP-1808.fasta | | | | |
| | Digestion | trypsin | trypsin | | trypsin | |
| | Max miss | 2 | 3 | | 2 | |
| | Fixed Modification | Carbamidomethyl on Cys | Carbamidomethyl on Cys | | Carbamidomethyl on Cys | |
| | Variable | Acetyl on Protein N- | Acetyl on Protein N-Term/Oxidation on | | Acetyl on Protein N-Term/Oxidation on Met | |
| **Modifications** | Mode | N-glycan | N-glycan | | N-Glycan | |
| | Glyco database | N-Human | 182 Human | | 182 Human | |
| **QC** | GPSM filter | 1%FDR | Score≥200&\|logProb\|≥2 | | 1%FDR | |
| **MS** | Fragmentation | HCD | HCD | | HCD | |
| | Precursor | 4ppm | 4ppm | | 20ppm | |
| | Fragment | 20ppm | 20ppm | | 20ppm | |
| **Quantification** | Quantification Software | pGlycoQuant | Byologic | pGlycoQuant | MSFragger-Glyco | pGlycoQuant |
| | Quant type | Label free | | | | |
| | Quant level | MS1 | MS1 | MS1 | MS1 | MS1 |
| | Precursor | 20ppm | — | 20ppm | 10ppm | 20ppm |
| | Fragment | — | | | | |
| | RT tolerance | 2min | — | 2min | — | 2min |

### 2-3 TMT-labeled mouse liver data

| | | | | | | |
|---|---|---|---|---|---|---|
| **Dataset** | TMT-labeled mouse liver data | | | | | |
| **Software** | Analysis Software | pGlyco3+pGlycoQuant | Byonic+Byologic | Byonic+pGlycoQuant | MSFragger-Glyco+ MSFragger-Glyco | MSFragger- Glyco+pGlycoQuant |
| | Identification Software | pGlyco3 | Byonic | | MSFragger-Glyco | |
| **Peptide** | Protein database | Human-H.sapiens-SP-1808.fasta | | | | |
| | Digestion | trypsin | trypsin | | trypsin | |
| | Max miss | 2 | 3 | | 2 | |
| | Fixed | Carbamidomethyl on | Carbamidomethyl on Cys | | Carbamidomethyl on Cys | |
| | Variable | Acetyl on Protein N- | Acetyl on Protein N-Term/Oxidation on | | Acetyl on Protein N-Term/Oxidation on Met | |
| **Glyco** | Mode | N-glycan | N-glycan | | N-Glycan | |
| | Glyco database | N-Human | 182 Human | | 182 Human | |
| **QC** | GPSM filter | 1%FDR | Score≥200&\|logProb\|≥2 | | 1%FDR | |
| **MS** | Fragmentation | HCD | HCD | | HCD | |
| | Precursor | 4ppm | 4ppm | | 20ppm | |
| | Fragment | 20ppm | 20ppm | | 20ppm | |
| **Quantification** | Quantification | pGlycoQuant | Byologic | pGlycoQuant | MSFragger-Glyco | pGlycoQuant |
| | Quant type | TMT | | | | |
| | Quant level | MS2 | — | MS2 | MS2 | MS2 |
| | Precursor | — | | | | |
| | Fragment | 20ppm | — | 20ppm | — | 20ppm |
| | RT tolerance | — | | | | |

### Supplementary Table 3 Quantitation results of all software tools

#### 3-1 SILAC-labeled 293T cell data

| **Dataset** | **SILAC-labeled 293T cell data** | | | | |
|---|---|---|---|---|---|
| **Identification** | Byonic | Byonic | MSFragger-Glyco | MSFragger-Glyco | pGylco3 |
| **Quantitation** | Byologic | pGlycoQuant | MSFragger-Glyco | pGlycoQuant | pGlycoQuant |
| **Number of PSM** | N/A | 3037 | N/A | 654 | 2295 |
| **PMVL(%)** | N/A | 1.05% | N/A | 2.60% | 0.17% |
| **PMVT(%)** | N/A | 0.66% | N/A | 1.30% | 0.09% |
| **Pearson** | N/A | 0.861 | N/A | 0.872 | 0.849 |
| **Standard Deviation** | N/A | 0.912 | N/A | 0.961 | 0.847 |
| **Number of Glycopeptide** | 985 | 738 | 361 | 309 | 597 |
| **PMVL(%)** | 53.40% | 1.22% | 11.08% | 0.971% | 0.34% |
| **PMVT(%)** | 26.75% | 0.68% | N/A | 0.485% | 0.17% |
| **Pearson** | 0.665 | 0.900 | N/A | 0.897 | 0.887 |
| **Standard Deviation** | 1.693 | 0.920 | 0.975 | 0.919 | 0.936 |
| **Number of Glycoprotein** | N/A | 209 | 148 | 146 | 204 |
| **PMVL(%)** | N/A | 1.44% | 4.73% | 0.068% | 0.98% |
| **PMVT(%)** | N/A | 0.96% | N/A | 0.034% | 0.49% |
| **Pearson** | N/A | 0.907 | N/A | 0.935 | 0.913 |
| **Standard Deviation** | N/A | 0.845 | 0.769 | 0.729 | 0.823 |

### 3-2 label-free Hela cell data

| Dataset | label-free Hela cell data | | | | |
|---|---|---|---|---|---|
| **Identification** | Byonic | Byonic | MSFragger-Glyco | MSFragger-Glyco | pGylco3 |
| **Quantitation** | Byologic | pGlycoQuant | MSFragger-Glyco | pGlycoQuant | pGlycoQuant |
| **Number of PSM** | N/A | 31491 | N/A | 7849 | 26288 |
| **PMVL(%)** | N/A | 0.87% | N/A | 0.15% | 0.01% |
| **PMVT(%)** | N/A | 0.40% | N/A | 0.04% | 0.000001% |
| **Pearson** | N/A | 0.980-0.992 | N/A | 0.980-0.993 | 0.984-0.995 |
| **Standard Deviation** | N/A | 0.144-0.317 | N/A | 0.148-0.312 | 0.133-0.300 |
| **Number of Glycopeptide** | 2716 | 2740 | 607 | 1651 | 2348 |
| **PMVL(%)** | 61.45% | 1.50% | 27.68% | 0.18% | 0% |
| **PMVT(%)** | 30.12% | 0.66% | 13.45% | 0.04% | 0% |
| **Pearson** | 0.940- | 0.985-0.994 | 0.940-0.987 | 0.985-0.994 | 0.988-0.996 |
| **Standard Deviation** | 0.186- | 0.190-0.357 | 0.159-0.393 | 0.184-0.320 | 0.174-0.330 |
| **Number of Glycoprotein** | N/A | 405 | 333 | 362 | 483 |
| **PMVL(%)** | N/A | 2.47% | 37.24% | 0.00% | 0% |
| **PMVT(%)** | N/A | 1.40% | 27.83% | 0.00% | 0% |
| **Pearson** | N/A | 0.991-0.998 | 0.971-0.998 | 0.992-0.997 | 0.994-0.998 |
| **Standard Deviation** | N/A | 0.20-0.286 | 0.068-0.244 | 0.136-0.265 | 0.132-0.285 |

### 3-3 TMT-labeled mouse liver data

| Dataset | TMT-labeled mouse liver data | | | | |
|---|---|---|---|---|---|
| **Identification** | Byonic | Byonic | MSFragger-Glyco | MSFragger-Glyco | pGylco3 |
| **Quantitation** | Byologic | pGlycoQuant | MSFragger-Glyco | pGlycoQuant | pGlycoQuant |
| **Number of PSM** | N/A | 252 | 73 | 73 | 128 |
| **PMVL(%)** | N/A | 2.78% | 5.48% | 5.48% | 4.69% |
| **PMVT(%)** | N/A | 2.18% | 4.79% | 4.79% | 3.52% |
| **Pearson** | N/A | 0.869 | 0.988 | 0.988 | 0.681 |
| **Standard Deviation** | N/A | 0.213 | 0.124 | 0.124 | 0.249 |
| **Number of Glycopeptide** | N/A | 151 | 53 | 64 | 100 |
| **PMVL(%)** | N/A | 1.99% | 3.77% | 3.13% | 2.00% |
| **PMVT(%)** | N/A | 1.66% | 1.89% | 3.13% | 1.50% |
| **Pearson** | N/A | 0.901 | 0.985 | 0.986 | 0.837 |
| **Standard Deviation** | N/A | 0.183 | 0.101 | 0.137 | 0.243 |
| **Number of Glycoprotein** | N/A | 50 | 42 | 55 | 52 |
| **PMVL(%)** | N/A | 2.00% | 4.76% | 1.82% | 0.00% |
| **PMVT(%)** | N/A | 2.00% | 2.38% | 1.82% | 0.00% |
| **Pearson** | N/A | 0.981 | 0.990 | 0.993 | 0.938 |
| **Standard Deviation** | N/A | 0.137 | 0.097 | 0.074 | 0.231 |

**Supplementary Figure 1 Workflow of pGlycoQuant.**

pGlycoQuant first reads the identification results, and then extracts the quantitation signals. Finally, the quantitation results are calculated.

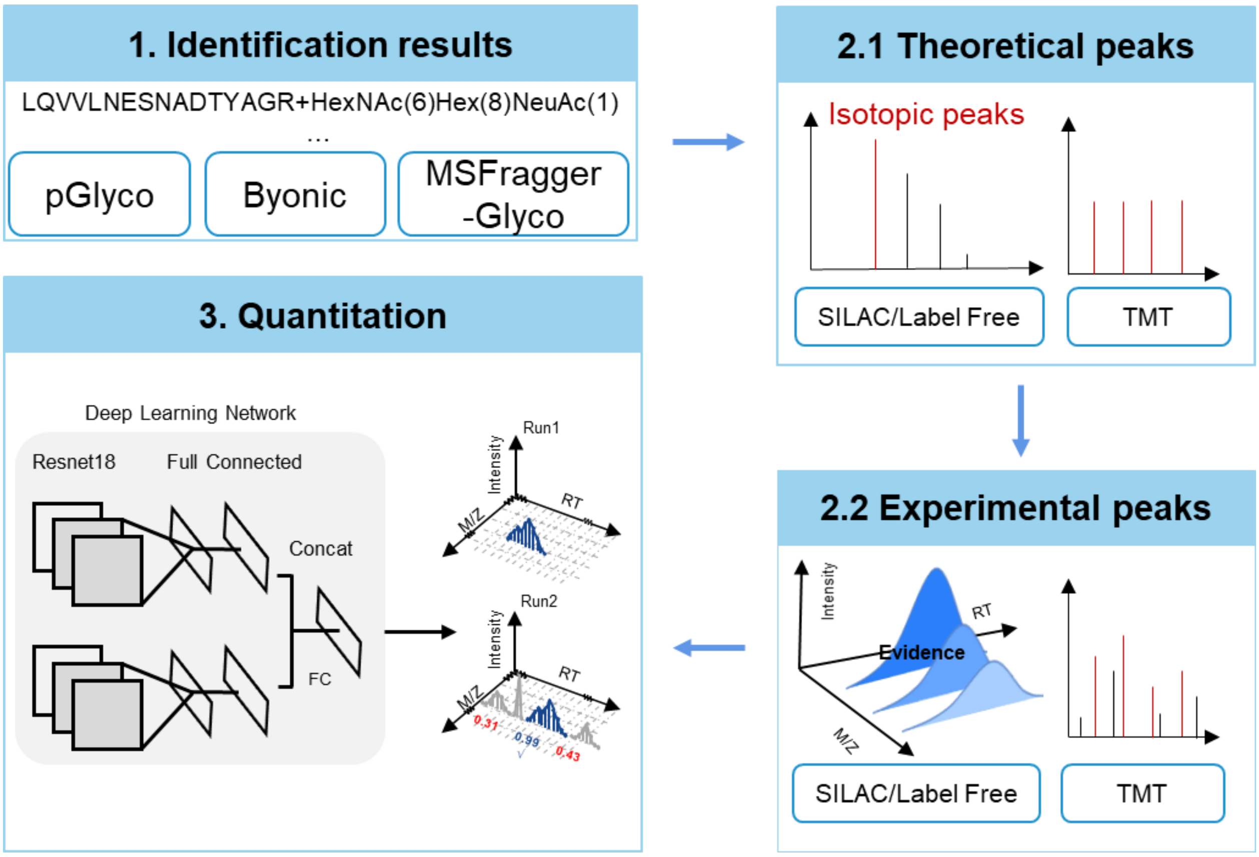

**Supplementary Figure 2 Deep-learning based evidence matching model.**

We consider the evidence as a matrix, similar to a picture in computational vision, so ResNet18 could be used to transform the matrix to a vector. Two fully connected (FC) network are used to calculate the similarity score of the given two evidence.

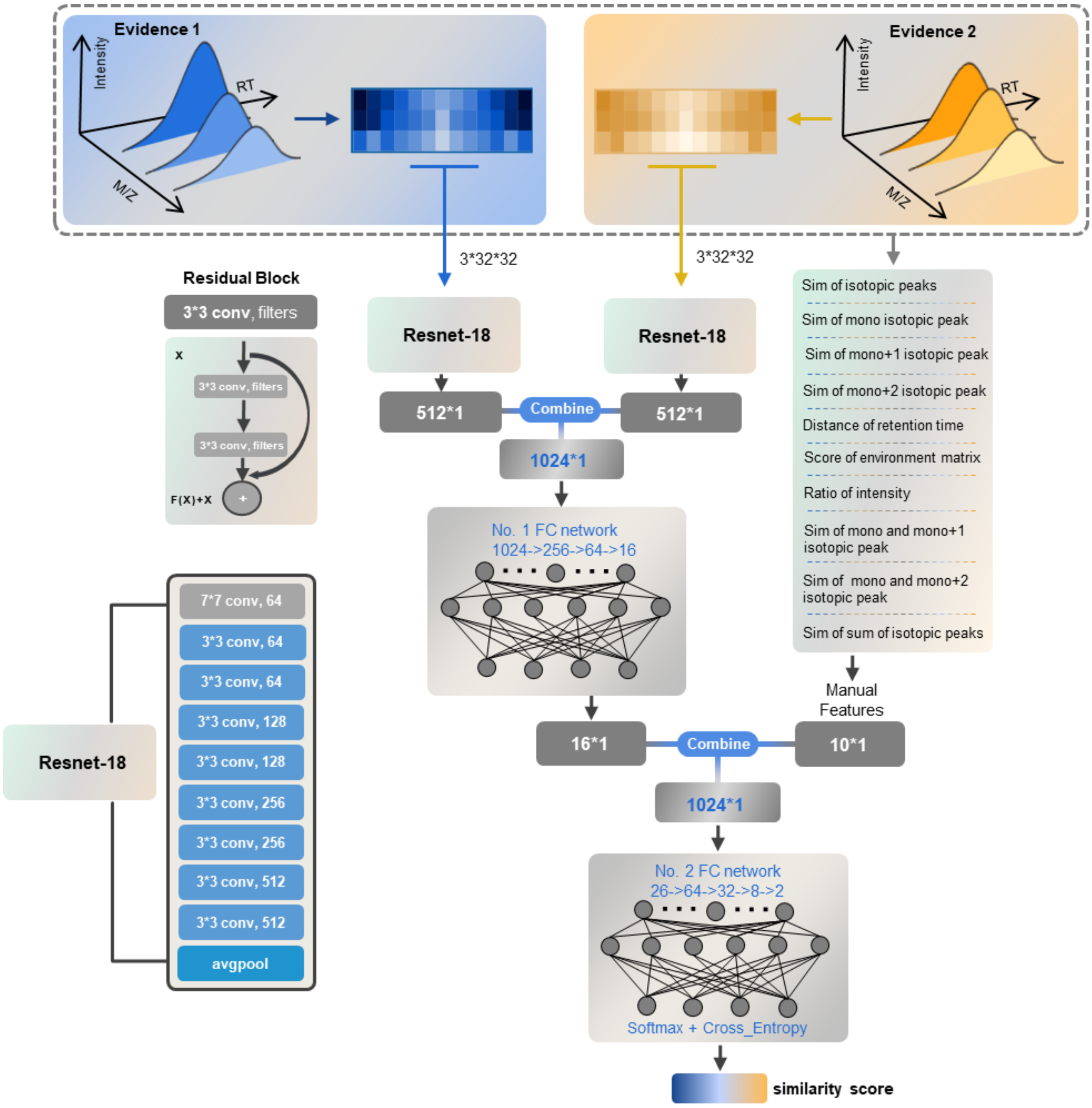

**Supplementary Figure 3 Scoring distribution of pGlycoQuant.**

We used the quantitative results of label-free data to illustrate scoring distribution of pGlycoQuant. The blue area represents the intensity ratio between 1/3 and 3 (the quantitative result is considered correct), the red area represents the result of the intensity ratio outside this range (the quantitative result is considered to be wrong). It can be seen that there are few results with a large ratio. To avoid the influence of potential wrong results on the subsequent bioinformatics analysis, we recommend select the results with matching probability scores greater than or equal to 0.6 (green dotted line).

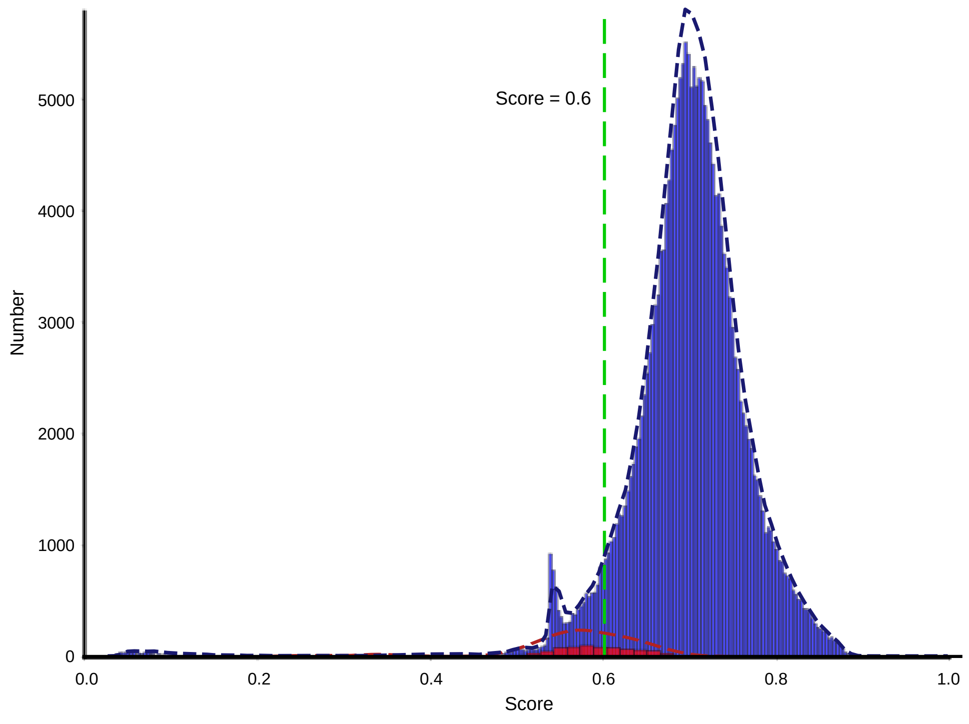

**Supplementary Figure 4 PMVT and PMVL.**

We define two indicators, proportion of missing value in line (PMVL) and proportion of missing value in total (PMVT). PMVL (%) is the number of peptides or proteins with more than one missing value among all samples divided by the number of all peptide or proteins. PMVT (%) is the number of individual missing value divided by the number of all quantitation values.

In this figure, a block indicates a quantitation value. Different blue color indicates different intensity, specifically, white color indicates missing value. Furthermore, we use red block to indicate the missing value number. The deeper the red color, the more missing values in line.

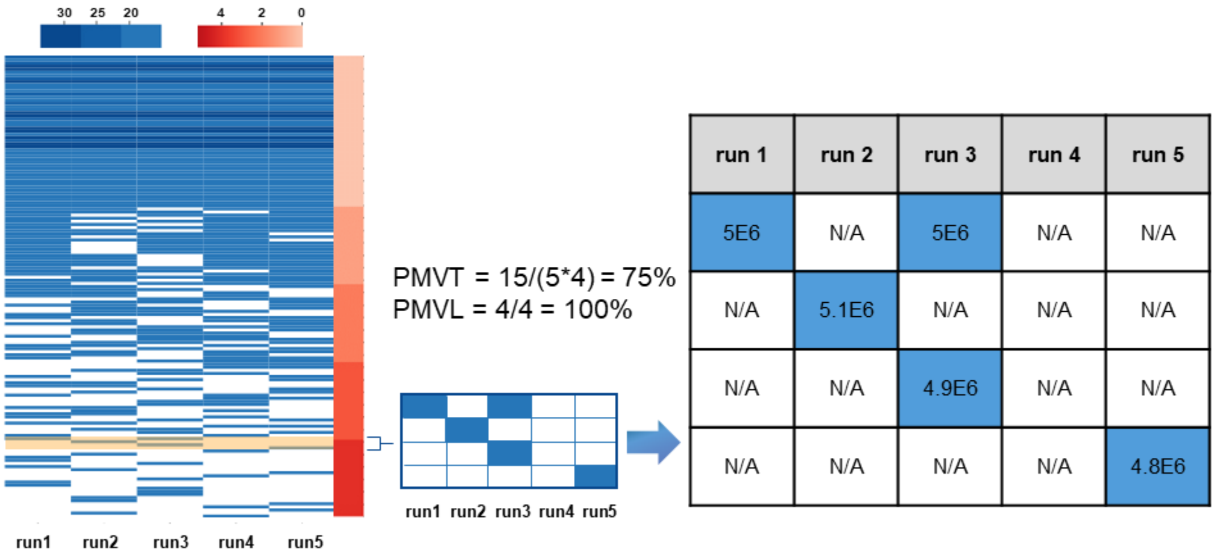

**Supplementary Figure 5 An example of missing values from label-free data.**

This figure illustrates the true signals of glycopeptide LAIMVNGSFK (Hex(9)HexNAc(2), 2+). These signals are existed and could be extracted by pGlycoQuant, but missed by other software tools.

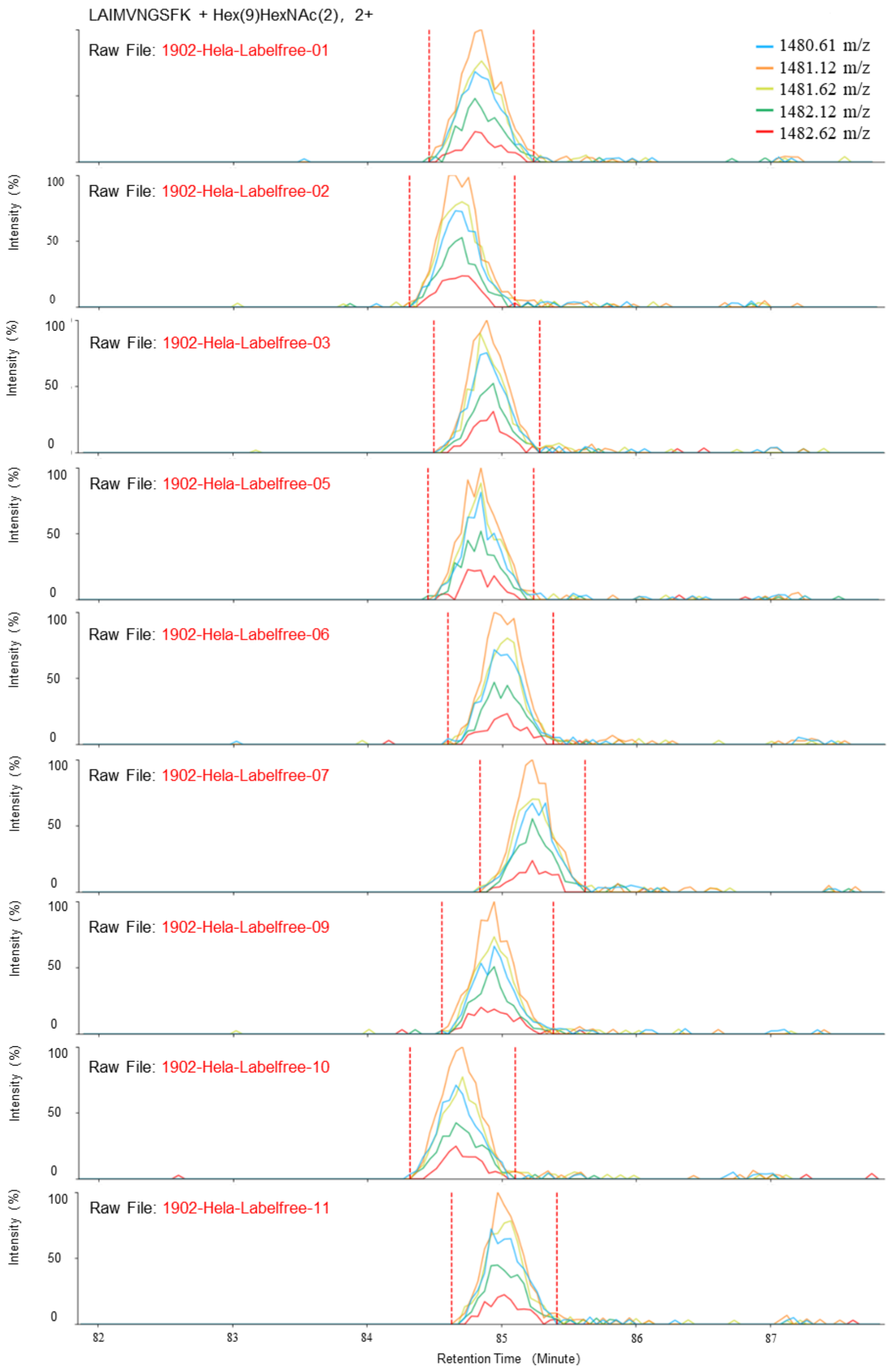

**Supplementary Figure 6 An example of missing values from SILAC data.**

This figure illustrates the true signals of glycopeptide EERPLNASALK (HexNAc(2)Hex(8), 3+). These signals are existed and could be extracted by pGlycoQuant, but missed by other software tools.

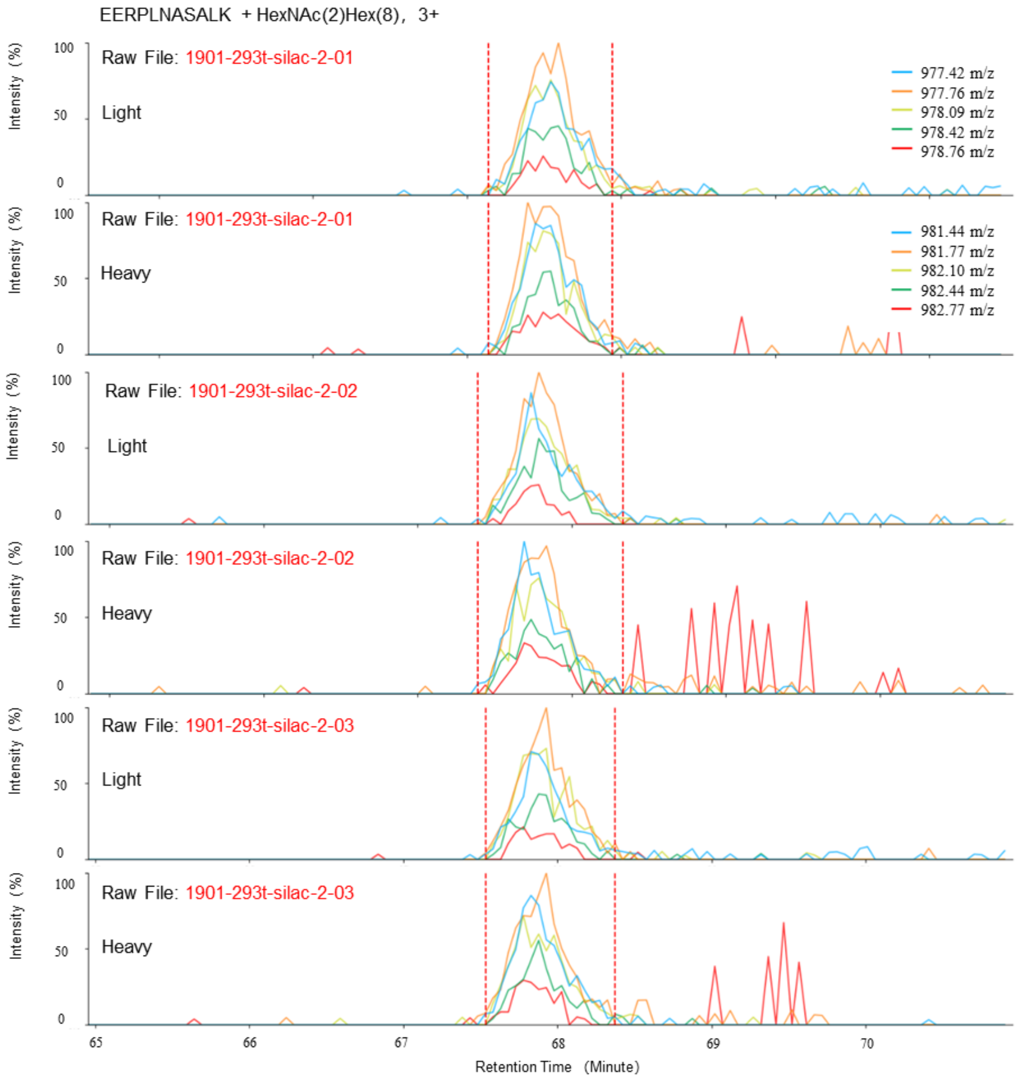

**Supplementary Figure 7 An example of missing values from TMT-labeled data.**

Compare to SILAC and label-free data, it is relatively easy to get the quantitation results for the TMT-labeled data, as only report ion should be considered. However, all software tools report a number of missing values, include pGlycoQuant. We check the true signals of the peptide, finding that no experimental peaks in the MS/MS scans.

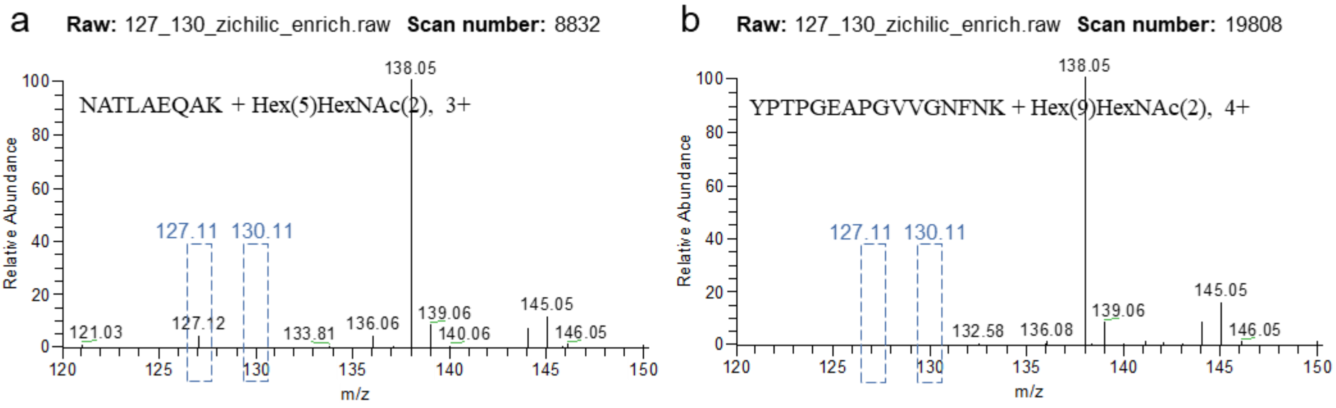

**Supplementary Figure 8 Quantitative results of SILAC data at peptide level.** (a) The correlation coefficient of quantitative intensity of each software. The lower left part is the scatter distribution diagram of intensity of light peptide and heavy peptide, and the upper right part is the calculated Pearson linear correlation coefficient. (b) The standard deviation of intensity ratio of each software. The lower left part is the violin diagram of intensity ratio of light peptide and heavy peptide, and the upper right part is the calculated standard deviation.

We evaluated the performance of quantitative results of each software on SILAC data and proved that pGlycoQuant can achieve better quantitative accuracy. From the following figure a, it can be seen that the quantitative results reported by pGlycoQuant have a higher correlation. It should be noted that MSFragger-Glyco's quantitative results from running SILAC data are the intensity ratios of light and heavy peptides, not the reported intensities, so there is no MSFragger-Glyco correlation result. Similarly, as shown in the following figure b the quantitative result reported by pGlycoQuant has a lower standard deviation.

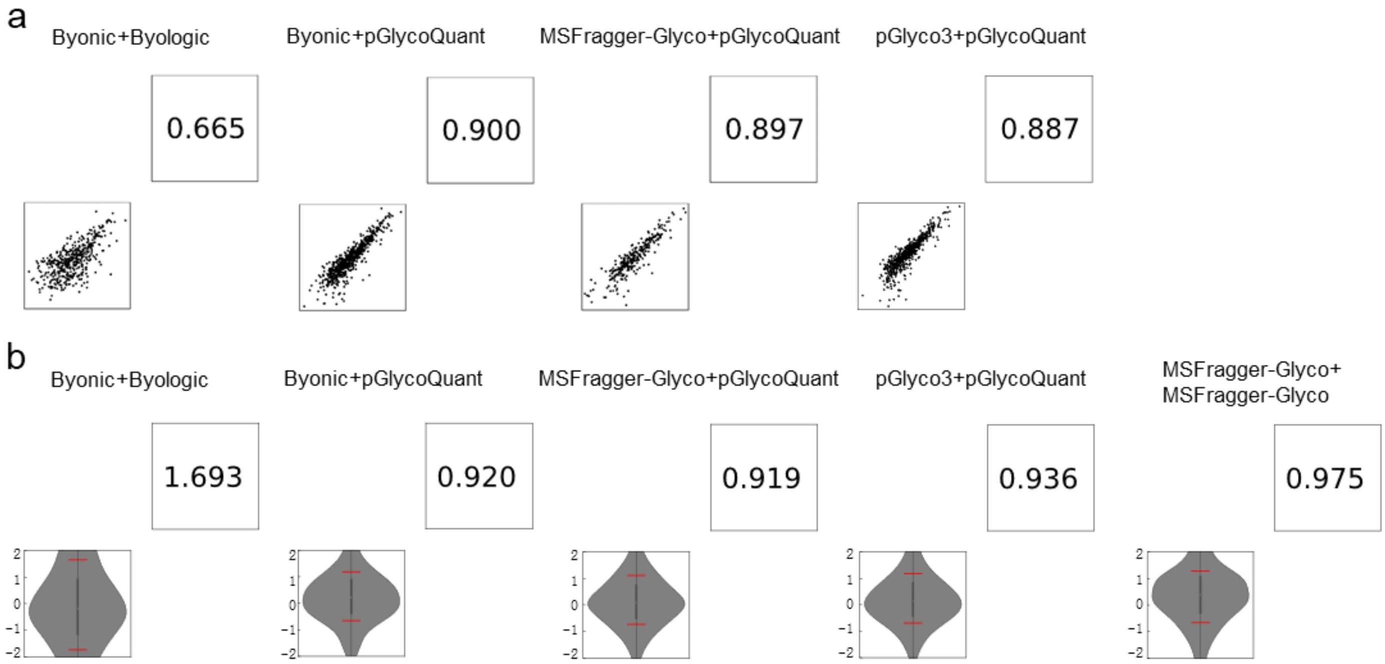

**Supplementary Figure 9 The performance of quantitative results of each software on label-free data**. (a) The correlation coefficient of quantitative intensity of each software. The lower left part is the scatter distribution diagram of intensity between two repeated data, and the upper right part is the calculated Pearson linear correlation coefficient. (b) The standard deviation of intensity ratio of each software. The lower left part is the violin diagram of intensity ratio between two repeated data, and the upper right part is the calculated standard deviation.

We evaluated the performance of quantitative results of each software on label-free data and proved that pGlycoQuant can achieve better quantitative accuracy. From the following figure a, it can be seen that the quantitative results reported by pGlycoQuant have a higher correlation. Similarly, the quantitative result reported by pGlycoQuant has a lower standard deviation.

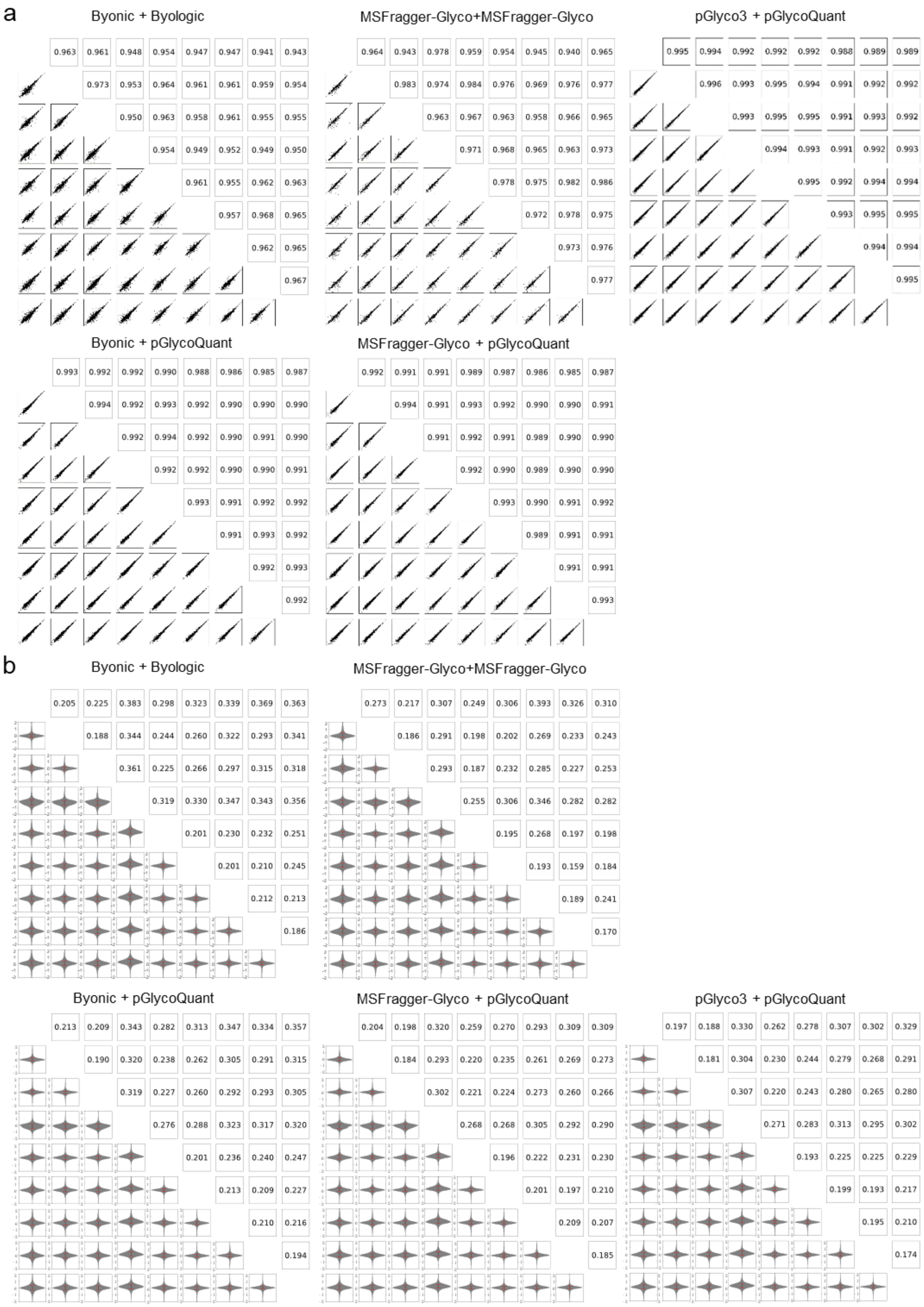
a
Byonic + Byologic
0.963 0.961 0.948 0.954 0.947 0.947 0.941 0.943
0.973 0.953 0.964 0.961 0.961 0.959 0.954
0.950 0.963 0.958 0.961 0.955 0.955
0.954 0.949 0.952 0.949 0.950
0.961 0.955 0.962 0.963
0.957 0.968 0.965
0.962 0.965
0.967
MSFragger-Glyco+MSFragger-Glyco
0.964 0.943 0.978 0.959 0.954 0.945 0.940 0.965
0.983 0.974 0.984 0.976 0.969 0.976 0.977
0.963 0.967 0.963 0.958 0.966 0.965
0.971 0.968 0.965 0.963 0.973
0.978 0.975 0.982 0.986
0.972 0.978 0.975
0.973 0.976
0.977
pGlyco3 + pGlycoQuant
0.995 0.994 0.992 0.992 0.992 0.988 0.989 0.989
0.996 0.993 0.995 0.994 0.991 0.992 0.992
0.993 0.995 0.995 0.991 0.993 0.992
0.994 0.993 0.991 0.992 0.993
0.995 0.992 0.994 0.994
0.993 0.995 0.995
0.994 0.994
0.995
Byonic + pGlycoQuant
0.993 0.992 0.992 0.990 0.988 0.986 0.985 0.987
0.994 0.992 0.993 0.992 0.990 0.990 0.990
0.992 0.994 0.992 0.990 0.991 0.990
0.992 0.992 0.990 0.990 0.991
0.993 0.991 0.992 0.992
0.991 0.993 0.992
0.992 0.993
0.992
MSFragger-Glyco + pGlycoQuant
0.992 0.991 0.991 0.989 0.987 0.986 0.985 0.987
0.994 0.991 0.993 0.992 0.990 0.990 0.991
0.991 0.992 0.991 0.989 0.990 0.990
0.992 0.990 0.989 0.990 0.990
0.993 0.990 0.991 0.992
0.989 0.991 0.991
0.991 0.991
0.993
b
Byonic + Byologic
0.205 0.225 0.383 0.298 0.323 0.339 0.369 0.363
0.188 0.344 0.244 0.260 0.322 0.293 0.341
0.361 0.225 0.266 0.297 0.315 0.318
0.319 0.330 0.347 0.343 0.356
0.201 0.230 0.232 0.251
0.201 0.210 0.245
0.212 0.213
0.186
MSFragger-Glyco+MSFragger-Glyco
0.273 0.217 0.307 0.249 0.306 0.393 0.326 0.310
0.186 0.291 0.198 0.202 0.269 0.233 0.243
0.293 0.187 0.232 0.285 0.227 0.253
0.255 0.306 0.346 0.282 0.282
0.195 0.268 0.197 0.198
0.193 0.159 0.184
0.189 0.241
0.170
Byonic + pGlycoQuant
0.213 0.209 0.343 0.282 0.313 0.347 0.334 0.357
0.190 0.320 0.238 0.262 0.305 0.291 0.315
0.319 0.227 0.260 0.292 0.293 0.305
0.276 0.288 0.323 0.317 0.320
0.201 0.236 0.240 0.247
0.213 0.209 0.227
0.210 0.216
0.194
MSFragger-Glyco + pGlycoQuant
0.204 0.198 0.320 0.259 0.270 0.293 0.309 0.309
0.184 0.293 0.220 0.235 0.261 0.269 0.273
0.302 0.221 0.224 0.273 0.260 0.266
0.268 0.268 0.305 0.292 0.290
0.196 0.222 0.231 0.230
0.201 0.197 0.210
0.209 0.207
0.185
pGlyco3 + pGlycoQuant
0.197 0.188 0.330 0.262 0.278 0.307 0.302 0.329
0.181 0.304 0.230 0.244 0.279 0.268 0.291
0.307 0.220 0.243 0.280 0.265 0.280
0.271 0.283 0.313 0.295 0.302
0.193 0.225 0.225 0.229
0.199 0.193 0.217
0.195 0.210
0.174

**Supplementary Figure 10 The performance of quantitative results of each software on TMT labeling data**. (a) The correlation coefficient of quantitative intensity of each software. The lower left part is the scatter distribution diagram of intensity of two labels, and the upper right part is the calculated Pearson linear correlation coefficient. (b) The standard deviation of intensity ratio of each software. The lower left part is the violin diagram of intensity ratio of two labels, and the upper right part is the calculated standard deviation.

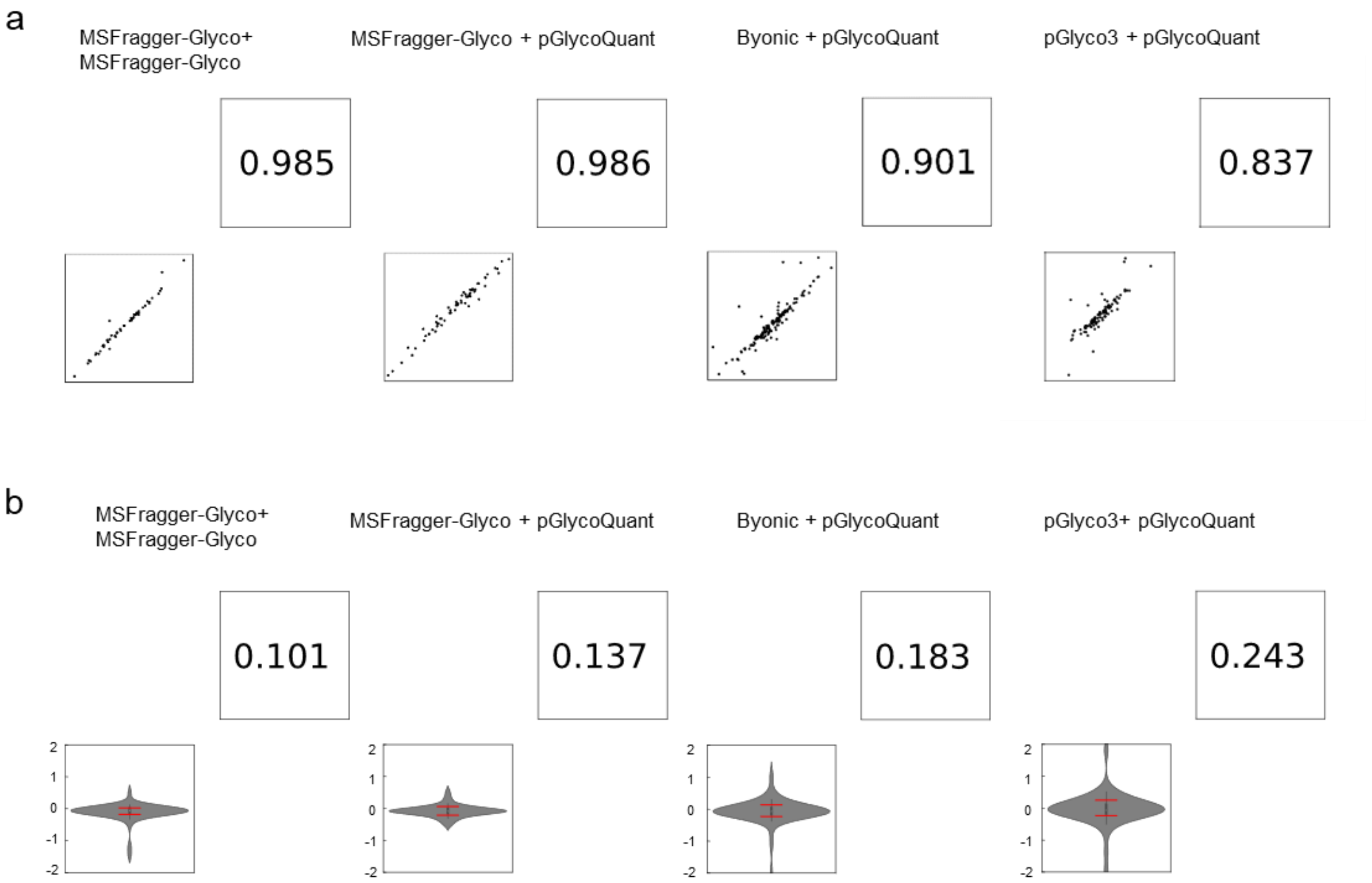

**Supplementary Figure 11 High repeatability of the proteome quantitative results among three cell lines**. (a) The normal distribution fitting line. (b) A correlation heat map of proteome quantitative ratio between different replicates.

a

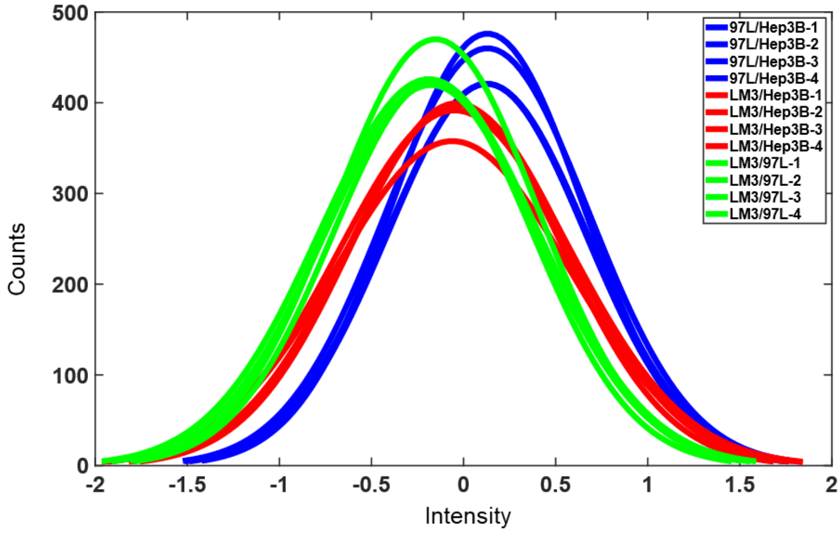

b

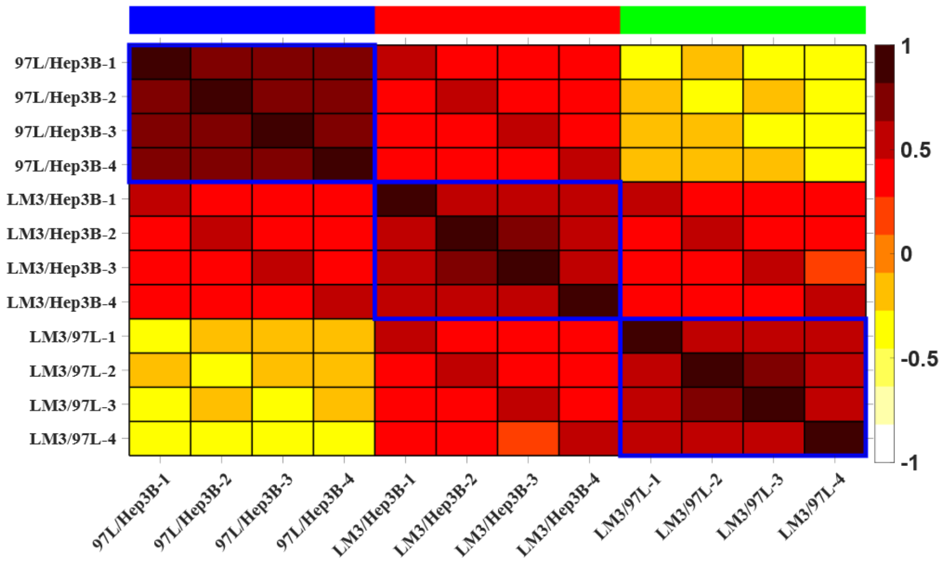

**Supplementary Figure 12 High repeatability of the intact glycopeptide quantitative results among three cell lines**. (a) The normal distribution fitting line. (b) A correlation heat map of intact glycopeptide quantitative ratio between different replicates.

a

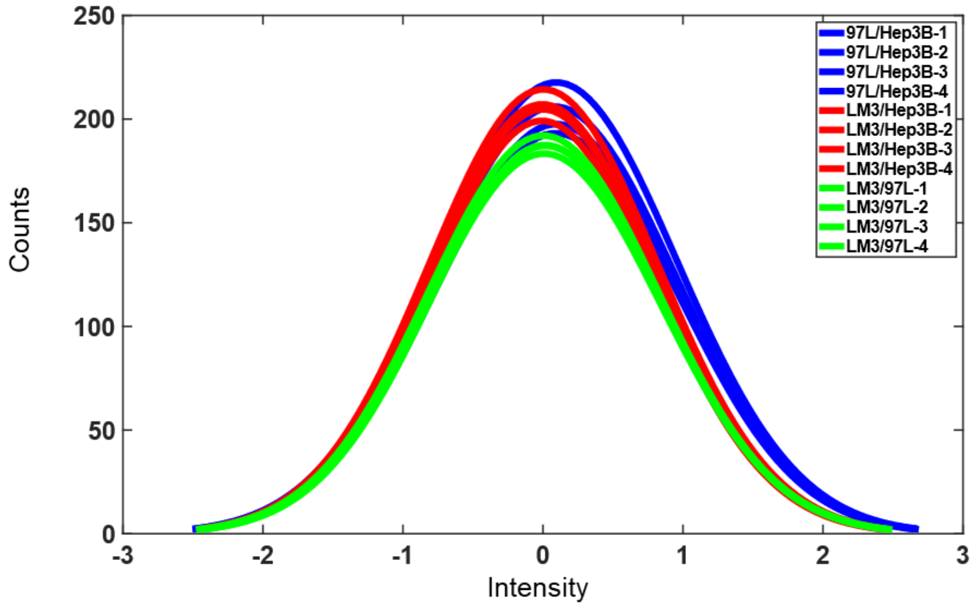

b

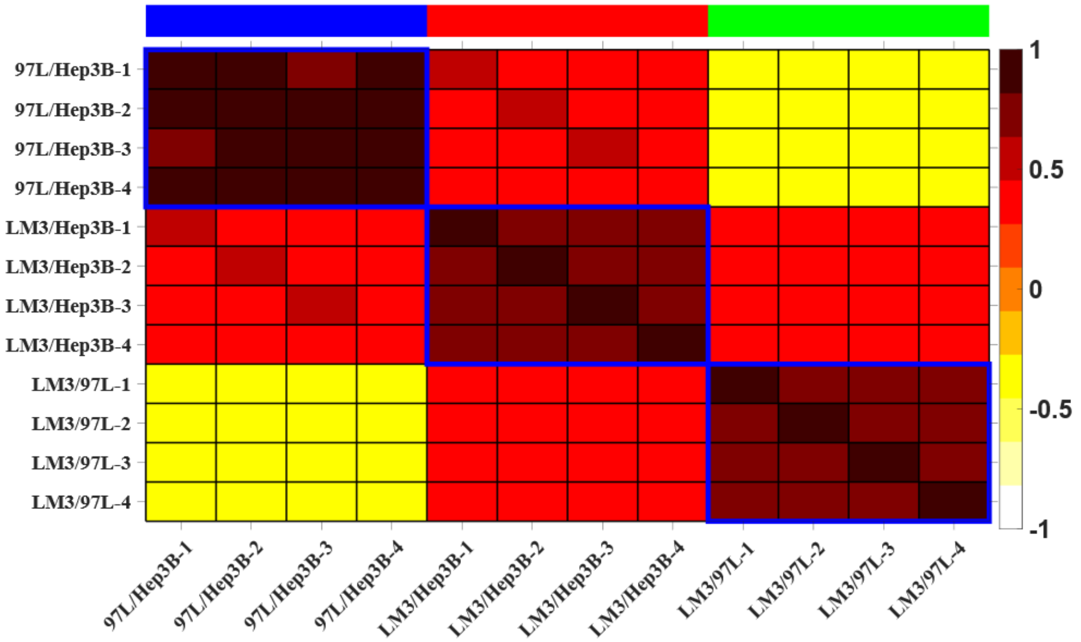

**Supplementary Figure 13 Gene ontology (GO) analyses of proteome** on cellular component (a), molecular function (b), and biological process (c).

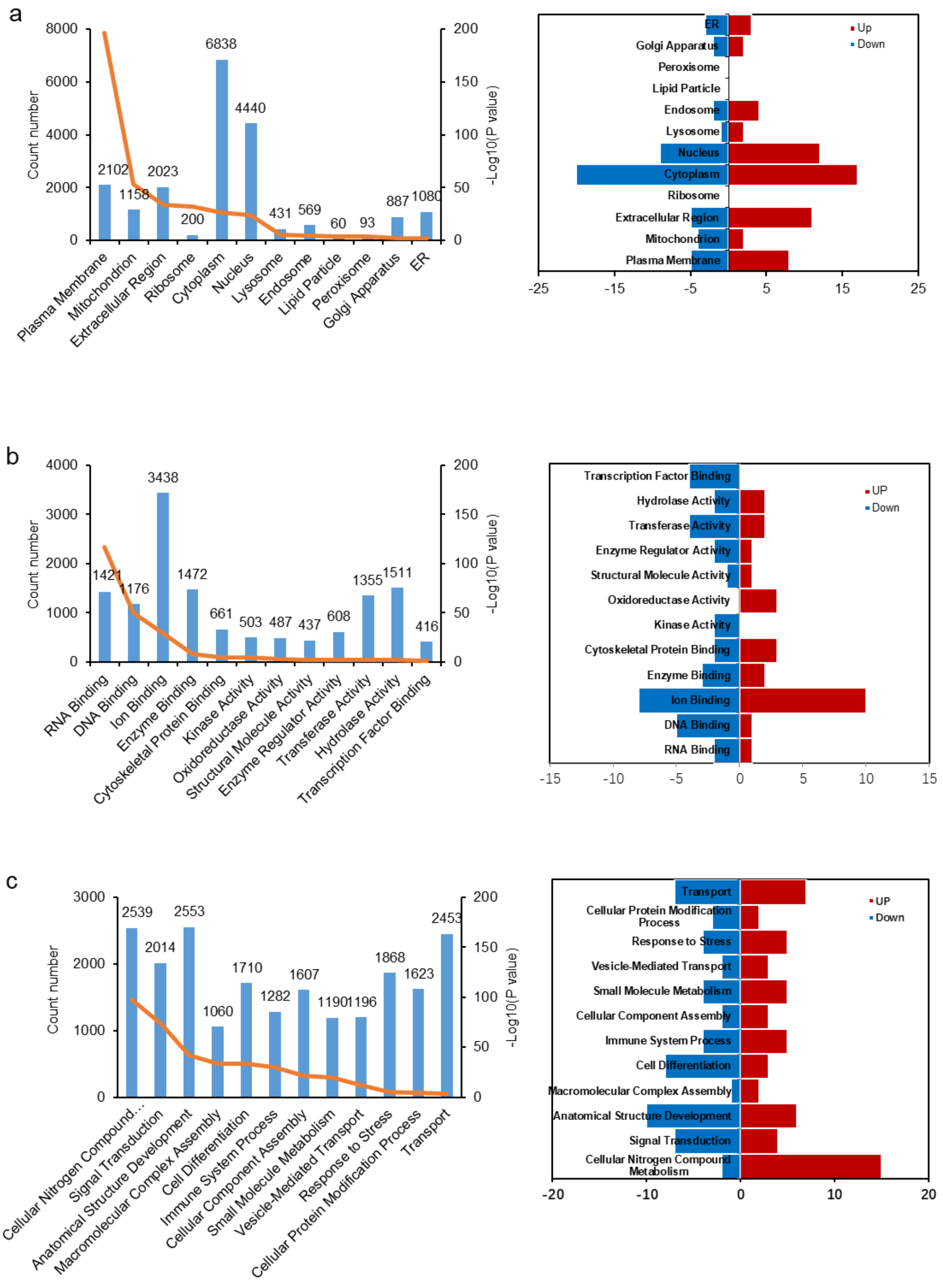

**Supplementary Figure 14 Gene ontology (GO) analyses of glycoproteome** on cellular component (a), molecular function (b), and biological process (c).

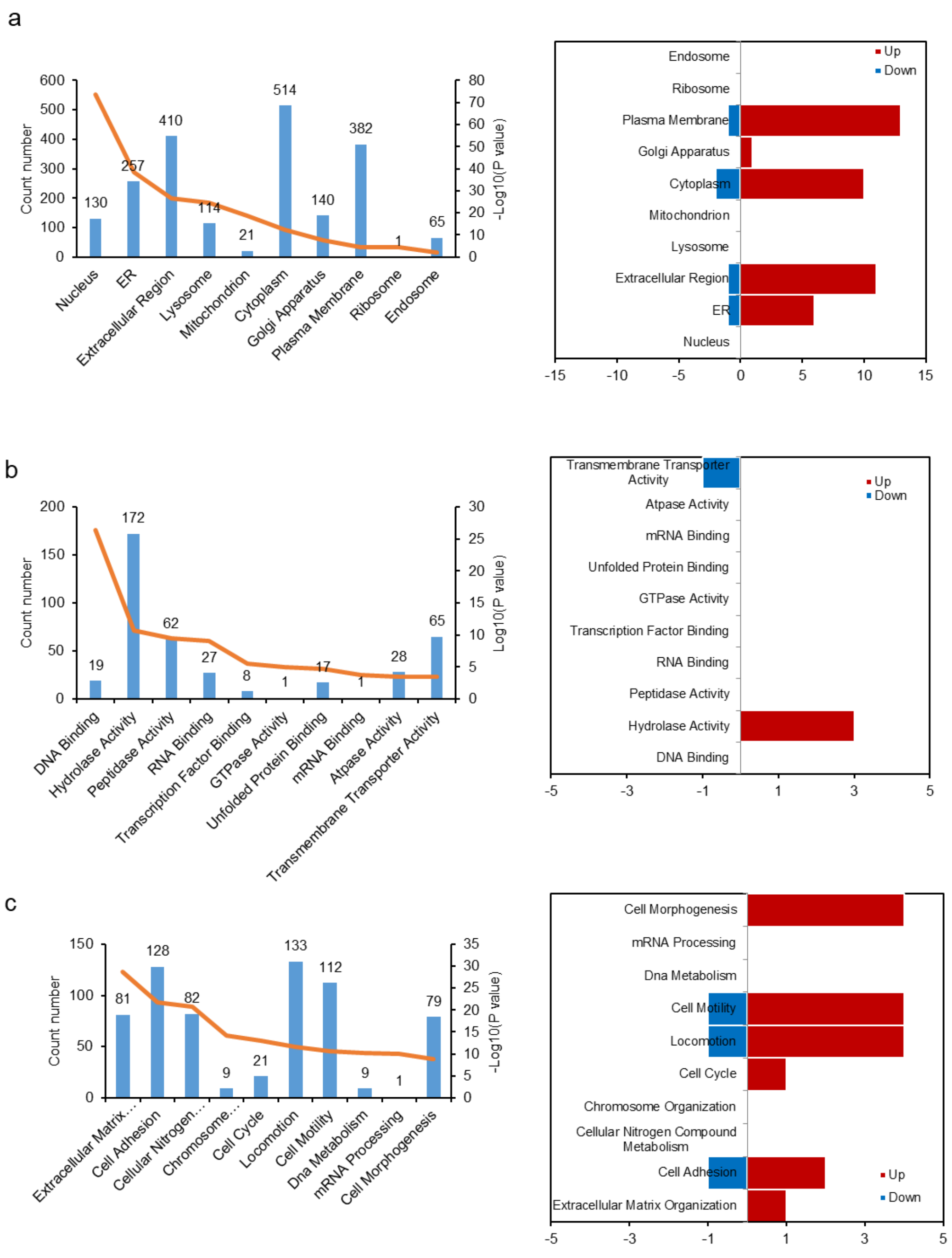

**Supplementary Figure 15 Distribution of glycan size and glycan type in three HCC cell lines**. (a) Distribution of glycan size (the number of saccharides units on a glycan) in different categories. (b) Distribution of different glycan types, including oligomannose, complex/hybrid, fucosylation, and sialylation in categories.

All is for all intact glycopeptides quantified in HCC cell lines. Up-up is for uniformly upregulated intact glycopeptides in three cell lines with increased metastatic potential. Down-down is for uniformly downregulated intact glycopeptides in three cell lines with increased metastatic potential.

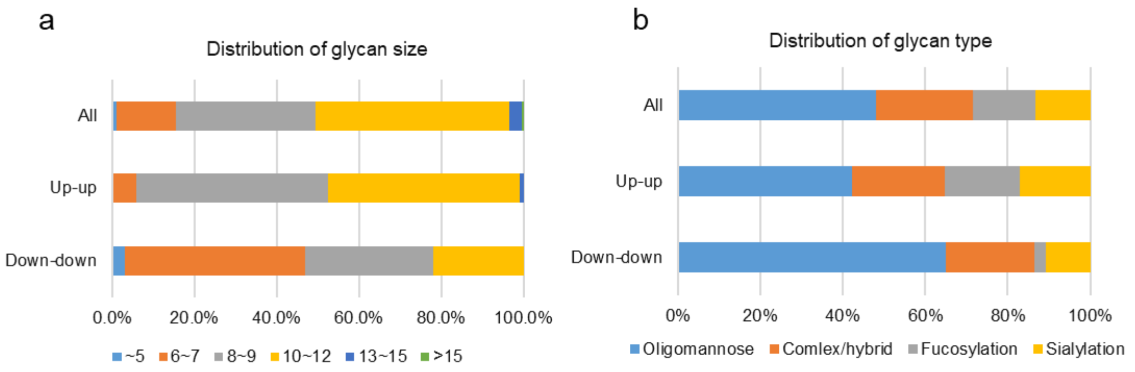

**Supplementary Figure 16 Differential expressed proteins of glycan-related genes in three HCC cell lines**. Blue fan represents human glycoside hydrolases. Orange fan represents human glycosyltransferases. Yellow fan represents other human glycan-related genes. Each glycan-related gene has two bars in the outermost circle. In the clockwise direction, the first bar represents the change of the protein in 97L/Hep3B, and the second bar represents the change of the protein in LM3/97L. Red bar stands for the upregulation. Blue bar stands for the downregulation. FUT8, which is yellow-highlighted, is a core-fucosylation transferase and is selected as the target glycosyltransferase in this study.

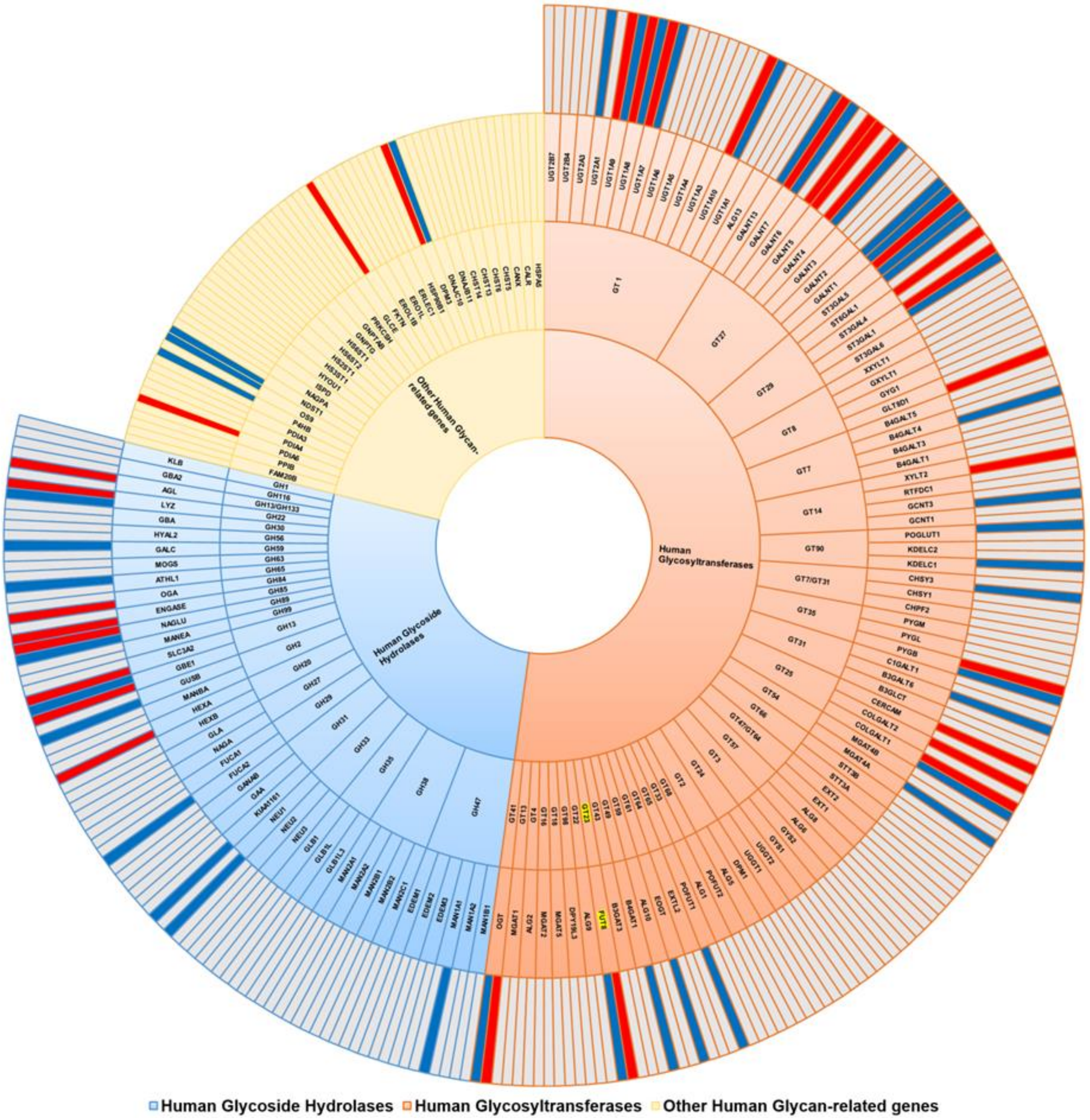

**Supplementary Figure 17 Site-specific glycosylation level of L1CAM in three HCC cell lines after normalization within a cell line or among three cell lines.** (a) is a table of the percentage of a site-specific glycan intensity to the total intensity of site-specific glycans in a cell line. (b) is a table of the percentage of a site-specific glycan intensity to the total intensity of site-specific glycans in three cell lines.

a

| Glycan(H,N,A,F) | N294 | | | N479 | | | N671 | | | N849 | | | N979 | | |
|---|---|---|---|---|---|---|---|---|---|---|---|---|---|---|---|
| | Hep3B | 97L | LM3 | Hep3B | 97L | LM3 | Hep3B | 97L | LM3 | Hep3B | 97L | LM3 | Hep3B | 97L | LM3 |
| 6 2 0 0 | | | | | | | | | | 0.514% | 5.385% | 5.111% | 0.018% | 0.032% | 0.039% |
| 7 2 0 0 | 0.054% | 0.058% | 0.109% | 3.067% | 6.121% | 6.100% | | | | 2.353% | 11.196% | 11.587% | 0.038% | 0.166% | 0.136% |
| 8 2 0 0 | 1.625% | 1.347% | | | | | 0.433% | 0.396% | 0.529% | 4.476% | 9.385% | 9.114% | 1.044% | 1.789% | 1.161% |
| 9 2 0 0 | 1.322% | 0.882% | 0.911% | | | | 0.280% | 0.178% | 0.174% | 0.299% | 0.685% | 0.491% | 1.724% | 2.302% | 1.175% |
| 10 2 0 0 | 0.069% | 0.042% | 0.051% | | | | | | | | | | | | |
| 5 3 0 1 | | | | | | | | | | | | | 0.278% | 0.527% | 0.031% |
| 5 3 0 2 | | | | | | | | | | | | | 0.337% | 0.468% | 0.107% |
| 5 3 1 0 | 0.104% | 0.045% | 0.075% | | | | | | | | | | 1.243% | 0.720% | 0.614% |
| 5 3 1 1 | | | | | | | | | | | | | 0.720% | 0.817% | 0.483% |
| 6 3 0 0 | | | | | | | | | | | | | 0.045% | 0.446% | 0.060% |
| 6 3 0 1 | | | | | | | | | | | | | 1.443% | 1.049% | 0.174% |
| 6 3 1 0 | 0.070% | 0.040% | 0.071% | | | | | | | | | | 1.546% | 0.946% | 0.725% |
| 8 3 0 0 | 0.040% | 0.026% | 0.076% | | | | | | | | | | | | |
| 5 4 0 0 | 0.123% | 0.159% | 0.072% | | | | | | | | | | | | |
| 5 4 0 1 | | | | | | | | | | | | | 0.437% | 0.494% | 0.293% |
| 5 4 0 2 | | | | | | | | | | | | | 1.699% | 3.060% | 3.018% |
| 5 4 1 0 | 5.684% | 1.727% | 2.536% | | | | | | | | | | 4.990% | 2.530% | 3.013% |
| 5 4 1 1 | 7.379% | 1.943% | 0.159% | | | | | | | | | | 52.462% | 41.619% | 45.721% |
| 5 4 1 2 | | | | | | | | | | | | | 0.120% | 0.482% | 0.116% |
| 5 4 2 0 | 3.761% | 2.543% | 4.218% | | | | | | | | | | 0.204% | 0.394% | 0.008% |

b

| Glycan(H,N,A,F) | N294 | | | N479 | | | N671 | | | N849 | | | N979 | | |
|---|---|---|---|---|---|---|---|---|---|---|---|---|---|---|---|
| | Hep3B | 97L | LM3 | Hep3B | 97L | LM3 | Hep3B | 97L | LM3 | Hep3B | 97L | LM3 | Hep3B | 97L | LM3 |
| 6 2 0 0 | | | | | | | | | | 0.015% | 0.564% | 4.427% | 0.001% | 0.003% | 0.034% |
| 7 2 0 0 | 0.002% | 0.006% | 0.094% | 0.089% | 0.641% | 5.283% | | | | 0.068% | 1.173% | 10.037% | 0.001% | 0.017% | 0.118% |
| 8 2 0 0 | 0.047% | 0.141% | 1.509% | | | | 0.013% | 0.041% | 0.458% | 0.130% | 0.983% | 7.894% | 0.030% | 0.187% | 1.005% |
| 9 2 0 0 | 0.038% | 0.092% | 0.789% | | | | 0.008% | 0.019% | 0.151% | 0.009% | 0.072% | 0.425% | 0.050% | 0.241% | 1.018% |
| 10 2 0 0 | 0.002% | 0.004% | 0.044% | | | | | | | | | | | | |
| 5 3 1 0 | 0.003% | 0.005% | 0.065% | | | | | | | | | | 0.036% | 0.075% | 0.532% |
| 6 3 1 0 | 0.002% | 0.004% | 0.062% | | | | | | | | | | 0.045% | 0.099% | 0.628% |
| 6 3 0 0 | | | | | | | | | | | | | 0.001% | 0.047% | 0.052% |
| 8 3 0 0 | 0.001% | 0.003% | 0.066% | | | | | | | | | | | | |
| 5 4 0 0 | 0.004% | 0.017% | 0.062% | | | | | | | | | | | | |
| 5 4 2 0 | 0.109% | 0.267% | 3.653% | | | | | | | | | | 0.006% | 0.041% | 0.007% |
| 5 4 1 0 | 0.165% | 0.181% | 2.197% | | | | | | | | | | 0.145% | 0.265% | 2.610% |
| 5 3 0 1 | | | | | | | | | | | | | 0.008% | 0.055% | 0.026% |
| 5 3 1 1 | | | | | | | | | | | | | 0.021% | 0.086% | 0.418% |
| 6 3 0 1 | | | | | | | | | | | | | 0.042% | 0.110% | 0.151% |
| 5 4 1 1 | 0.214% | 0.204% | 0.138% | | | | | | | | | | 1.523% | 4.361% | 39.603% |
| 5 4 0 1 | | | | | | | | | | | | | 0.013% | 0.052% | 0.254% |
| 5 3 0 2 | | | | | | | | | | | | | 0.010% | 0.049% | 0.093% |
| 5 4 0 2 | | | | | | | | | | | | | 0.049% | 0.321% | 2.614% |
| 5 4 1 2 | | | | | | | | | | | | | 0.003% | 0.050% | 0.101% |

**Supplementary Figure 18 Silencing L1CAM inhibited migration and invasion ability of LM3 cells**. (a) Western blot of L1CAM levels in 97L cells transfected with negative control (Ctrl) and L1CAM siRNA. (b) Wound healing assay in monolayers of knocked down LM3 cells (siCtrl and siL1CAM). Scratch area of the cells were detected with an inverted microscope (10X). (c) Transwell migration assay (upper) and (d) matrigel invasion assay (down) (Scale bar = 100 μm) in knocked down LM3 cells (siCtrl and siL1CAM).

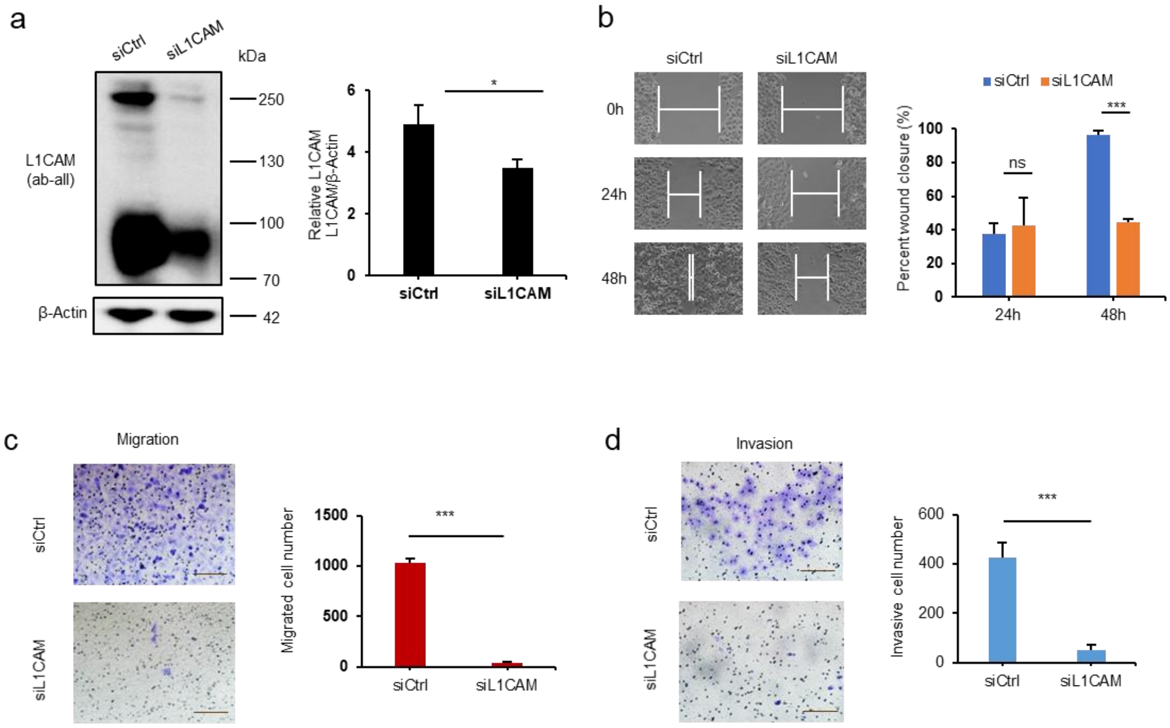

**Supplementary Note 1 Methods for sample preparation and LC-MS/MS**

**1.1 SILAC labeling of 293T sample preparation**

**Cell culture and labeling**

The K0R0 medium containing Dulbecco's modified Eagle's medium (DMEM, HyClone, Logan, UT, USA) supplemented with 10% dialyzed fetal bovine serum (FBS, Gibco, Grand Island, NY, USA) was prepared as light medium for light labeling. The K6R6 medium containing DMEM with L-lysine ($^{13}C_6$, 99%) and L-arginine ($^{13}C_6$, 99%) and 10% dialyzed FBS was prepared as heavy medium (K6R6-medium) for heavy labeling.

Split a dish of 293T cells into two separate dishes, each containing K0R0 medium and K6R6 medium respectively. The cells were cultured at 37°C in a humidified 5% CO2 incubator. When the cells are >80% confluent, rinse the cells with PBS, detach the cells with trypsin/EDTA and split the cells onto new dishes. Cell lines were grown for eight cell divisions in either K0R0 medium or K6R6 medium.

**Cell collection and protein extraction**

After SILAC labeling is complete, remove medium from the dishes, wash twice with ice cold PBS, and then completely remove PBS. The cells were collected in a tube and lysed in five-volume lysis buffer (4% SDS, 0.1M Tris/HCl, pH 8.0) with protease inhibitor (1mM PMSF, 1mM cocktail), followed by boiling at 100°C for 10 minutes, ultrasonication for 20 minutes and centrifugation at 12,000g at 18°C for 30 minutes to collect protein extracts. The protein concentration was determined by BCA method.

**Protein digestion**

The proteins extracted from K0R0-labeled cells and K6R6-labeled cells were mixed with 1:1. Then, the protein mixtures were reduced in 10mM dithiothreitol at 37 °C for 1 hour, and then alkylated in dark by 20mM iodoacetamide at room temperature for 30 minutes. After that, six volumes of precooled acetone were added to precipitate the proteins at −20°C for at least 3 hours. The precipitates were dissolved in a denaturing buffer (8M urea in 50mM $NH_4HCO_3$) following a ten-fold dilution with 50mM $NH_4HCO_3$. Trypsin was added to a final enzyme-tosubstrate ratio of 1:50 (w:w) and incubated at 37°C overnight. The reactions were terminated by adding trifluoroacetic acid with final concentration of 0.5%. Finally, all digests were centrifuged at 16,000 × g for 10 min and the supernatants were desalted using Sep-Pak C18 cartridges (Waters, USA). The desalted peptides were then lyophilized for subsequent enrichment procedures.

**1.2 Label free of Hela sample preparation**

Hela cells were cultured in DMEM supplemented with 10% dialyzed FBS at 37 °C in a humidified 5% CO2 incubator. The cells were collected in a tube, and washed twice with ice cold PBS. Proteins were then extracted from cells, digested and desalted following the same procedures in 293T sample preparation. The desalted peptides were lyophilized and stored at -80℃ for subsequent enrichment procedures.

**1.3 TMT labeling of mouse live sample preparation**

Mouse livers used in this study were dissected from mouse strain C57BL/6, male, aged 3 months. Three mice were anesthetized with avertin and killed. Livers were taken out after perfusion with 0.9% NaCl. The procedures were incompliance with ethical regulations and were approved by the ethics committee of Fudan University. The tissues were grinded with liquid nitrogen, collected in a tube and lysed in five-volume lysis buffer (4% SDS, 0.1M Tris/HCl, pH 8.0) with protease inhibitor (1mM PMSF, 1mM cocktail), followed by boiling at 100°C for 10 minutes, ultrasonication for 20 minutes and centrifugation at 12,000g at 18°C for 30 minutes to collect protein extracts. Proteins were digested and desalted following the same procedures for 293T sample preparation.

The digests were divided into two aliquots, each of which were labeled with the TMT6plex™ label reagents $TMT^6$-127 and $TMT^6$-130 following the TMT6plex™ isobaric label reagent product manual (Thermo Fisher Scientific, Waltham, MA, U.S.A.). Briefly, per 100μg protein digests were resuspended with 100μL of 100mM TEAB, pH 8.5. A total of 41μL of anhydrous acetonitrile was added into each 0.8mg vials and allowed to dissolve for 5 minutes with occasional vortex. After briefly centrifugation the tube, the 41μL of the reagent solution was added to each 100μL sample (containing 100μg protein digests) and incubated for 1 hour at room temperature. Then, 8μL of 5% hydroxylamine was added to the sample and incubated for 15 minutes to quench the reaction. Equal amounts of each samples were combined in a new microcentrifuge tube and lyophilized. Then the samples were cleaned up using Sep-Pak C18 cartridges (Waters, USA) and lyophilized for subsequent enrichment procedures

**1.4 SILAC labeling of HCC cell lines sample preparation**

Three hepatocellular carcinoma cell lines, Hep3B, MHCC97L and MHCCLM3, were adopted for SILAC labeling. The K0R0 medium containing DMEM supplemented with 10% dialyzed FBS was used as light medium for labeling of 97L. The K4R6 medium containing DMEM with L-lysine ($D_4$, 99%) and L-arginine ($^{13}C_6$, 99%) supplemented with 10% dialyzed FBS was used as median medium for labeling of Hep3B. The K8R10 medium containing DMEM with L-lysine ($^{13}C_6$, 99%; $^{15}N_2$, 99%) and L-arginine ($^{13}C_6$, 99%; $^{15}N_4$, 99%) supplemented with 10% dialyzed FBS was used as heavy medium for labeling of LM3. Hep3B, MHCC97L and MHCCLM3 cells were cultured in the above medium respectively following the same experimental procedures with that for SILAC-293T cells. After complete SILAC labeling, the three kind of cells were collected followed by protein digestion as that for 293T cells, respectively. Then the proteins extracted from the three cell lines were mixed with 1:1:1, digested, desalted using Sep-Pak C18 cartridges, and lyophilized for subsequent processing.

The digests were fractionated by hydrophilic interaction liquid chromatography (HILIC) using a Waters UPLC system coupled with a HILIC column (Welch, Ultimate HILIC Amide, 4.6× 250 mm, 5 μm). The digests were re-dissolved in phase A (10 mmol/L ammonium formate, $H_2O$, pH 4.5 adjusted with FA). The gradient was as follows: 100% B (75% ACN and 10 mmol/L ammonium formate) for 5 min, 100–90% B in 3 min, 90–40% B in 56 min, 40–20% B in 3 min, return

100% B in 0.1min, and hold until the end of gradient. The flow rate was 1 mL/min and the column temperature maintained at 45°C. The collection started at the 4$^{th}$ minute to the 70$^{th}$ minute with 1 minutes in turn and were combined as the following conditions: fraction 1 (1-11), fraction 2 (12-19), fraction 3 (20-27), fraction 4 (28-35), fraction 5 (36-43), fraction 6 (44-51), and fraction 8 (52-59), and fraction 9 (60-67). Then, each of fractions was desalted and divided into two parts. One part is used for LC-MS/MS analysis for proteome identification and quantitation. The other part is used for glycopeptide enrichment and LC-MS/MS analysis for intact glycopeptide identification and quantitation.

**1.5 Glycopeptide enrichment**

Glycopeptides were enriched by zwitterionic hydrophilic interaction liquid chromatography (ZIC-HILIC) method. Briefly, the desalted peptides of 500μg-1 mg were resuspended in 300 μL loading buffer containing 80% ACN and 1% TFA and then loaded onto a homemade micro-column containing 50 mg of ZIC-HILIC particles (Merck Millipore, Darmstadt, Germany) packed onto a C8 disk. The flow through was collected and reloaded onto the column for additional four times. Then, the column was washed with 200 μL loading buffer for four times, and finally eluted with 140 μL 0.1% TFA. The elution was collected and lyophilized.

**1.6 LC-MS/MS analysis for proteome**

All LC-MS/MS analyses were performed on LC-MS/MS on an Orbitrap Fusion Tribrid system (Thermo Fisher Scientific, Waltham, MA, USA) equipped with an EASY-nLC TM1100 system (Thermo Fisher Scientific, Waltham, MA, USA) that included a reverse-phase analytical column without the trap column. Solvent A was a 0.1% FA aqueous solution. Solvent B was ACN containing 0.1% FA. Detailed LC-MS/MS parameters are listed below.

| | | SILAC-labeled 293T cell | label-free HeLa cell | TMT-labeled mouse liver | SILAC-labeled HCC cell lines for intact glycopeptide | SILAC-labeled HCC cell lines for proteome |
|---|---|---|---|---|---|---|
| LC | Column | C18 column 50 cm×75 μm i.d. (Thermo) | | | | |
| | Flow rate | 250 nl/min | 250 nl/min | 200 nl/min | 200 nl/min | 200 nl/min |
| | total gradient time | 360 min | 360 min | 180 min | 240 min | 120 min |
| | Elute gradient | 1–30% B in 330 min, 30–45% B in 15 min, 45-90% in 1 min, 90% for 7 min, 90%-1% in 10 s, 1% for 6'50s | 1–30% B in 330 min, 30–45% B in 15 min, 45-90% in 1 min, 90% for 7 min, 90%-1% in 10 s, 1% for 6'50s | 5–35% B in 165 min, 35–50% B in 7 min, 50-90% in 1 min, 90% for 3 min, 90%-5% in 10 s, 5% for 3'50s | 1–20% B in 180 min, 20–30% B in 42 min, 30-90% in 3 min, 90% for 7 min, 90%-1% in 10 s, 1% for 7'50s | 3–8% B in 4 min, 8–24% B in 80 min, 24-35% in 24 min, 35-90% in 2 min, 90% for 4 min, 90%-3% in 10 s, 3% for 5'50s |
| MS1 | Scan range (m/z) | 350-2000 | 350-2000 | 350-2000 | 350-2000 | 350-1550 |
| | Resolution | 120,000 | | | | 240,000 |
| | Included charge state | 2-6 | | | | |
| | dynamic exclusion after n times | 1 | | | | 1 |
| | dynamic exclusion duration | 15 s | | | | 60 s |
| | Data acquisition mode | DDA | | | | |
| MS/MS | Resolution | 15,000 | | | | 30,000 |
| | Maximum injection time | 250 ms | 250 ms | 200 ms | 250 ms | 64 ms |
| | Collision energy | HCD@30%±10% | | | | HCD@36% |

**Supplementary Note 2 Methods for invitro molecular biology experiments**

**Cell lines and cell culture.** Hepatocellular carcinoma MHCC97L and MHCCLM3 cell lines were cultured in Dulbecco's modified Eagle's medium (DMEM, HyClone, Logan, UT, USA) supplemented with 10% fetal bovine serum (FBS, Gibco, Grand Island, NY, USA) and 10 U/mL penicillin streptomycin (Gibco by Invitrogen, Carlsbad, CA, USA) at 37 °C in a humidified 5% CO2 incubator.

**Western blot assay.** The cell lysate was collected and prepared in ice-cold RIPA lysis and extraction Buffer (Thermo Fisher Scientific, Waltham, MA, USA), supplemented with a complete protease inhibitor mixture (Roche Diagnostics, Penzberg, Germany). Total cellular proteins were resolved by SDS-PAGE and transferred to a polyvinylidene difluoride (PVDF) membrane. After blocking nonspecific binding with TBS/T (0.1%) containing 5% non-fat milk for 1 h at room temperature, the membrane was incubated with the following different primary antibodies: anti-FUT8 (66118-1-Ig, 1:1000), anti-FLAG (80010-1-RR, 1:5000), anti-β-Actin (66009-1-Ig, 1:10000) (Proteintech, Rosemont, IL, USA), anti-L1CAM ab-all (ab182407, 1:1000), anti-L1CAM ab3 (ab24345, 1:1000) and anti-β-Tubulin (ab151318, 1:1000) (Abcam, Cambridge, UK). The membrane was washed with TBS/T four times to remove the unbound antibody and then incubated with the secondary antibody (HRP-conjugated goat anti-mouse IgG or goat anti-rabbit IgG, 1: 10000; Affinity Bioscience, Jiangsu, China) for 1 h at room temperature. Protein bands were visualized with an ECL kit (BeyoECL Plus, Beyotime Institute of Biotechnology, Shanghai, China).

**Lectin enrichment and immunoblot assay.** A total of 1000 μg of cell lysate was mixed with 320 μL of agarose bound Pisum Sativum Agglutinin (PSA) lectin or agarose bound Lens Culinaris Agglutinin (LCA) lectin (Vector laboratories, Burlingame, CA, USA) in 650 μL binding buffer containing 2 mM $MnCl_2$, 2 mM $CaCl_2$, and 1 mM $NaCl_2$ and incubated with rotation at 4 °C overnight. Then, the beads were washed three times with binding buffer and subsequently extracted with SDS-PAGE sample buffer at 99 °C for 10 min. The samples were separated by 4-20% Bis-Tris-PAGE and subjected to immunoblotting with antibody anti-L1CAM (ab-all) (ab182407, 1:1000). For lectin blot, after separation through Bis-Tris—PAGE, the samples were transferred to PVDF membranes. The membrane was washed with TBS/T four times to remove the unbound antibody and then incubated with the secondary antibody (HRP-conjugated goat anti-mouse IgG or goat anti-rabbit IgG, 1: 10000; Affinity Bioscience, Jiangsu, China) for 1 h at room temperature. Protein bands were visualized with an ECL kit (BeyoECL Plus, Beyotime Institute of Biotechnology, Shanghai, China).

**Wound healing assay.** Cells were seeded into a 24-well culture plate with the number of $1 \times 10^5$ cells/well. When the cell density reached 90% confluency, a cell monolayer was scratched gently with a sterile pipette tip. After washing with PBS twice, fresh medium was added. Cell migration was observed under a microscope (DMi8, Leica, Wetzlar, Germany) and imaging was performed at different time points.

**Transwell migration assay.** Transwell assay was conducted using a transwell chamber (Corning, NY, USA) with an aperture of 8 μm to estimate cell migration. A total of $5 \times 10^4$ cells was suspended in 100 μL serum-free medium and then added into the upper chamber, whereas 600 μL of 10% FBS medium was added in the lower chamber. After incubation at 37 °C with 5% $CO_2$ for 36 h, the non-migrating cells on the upper chamber were removed with cotton swabs, and the migrated cells were fixed with methanol and stained with 0.1% crystal violet. Five fields were randomly selected and the number of migrated cells was counted under the microscope.

**Matrigel invasion assay.** As for the transwell invasion assay, the upper chamber membranes were coated with matrigel (Corning, NY, USA). $1 \times 10^5$ cells were added into the upper chamber in serum-free medium, the rest was the same as the migration assay.

**siRNA and plasmid transfection.** FUT8-specific siRNAs (sequence: 5′-GUGGAGUGAUCCUGGAUAUTT-3′, 5′-AUAUCCAGGAUCACUCCACTT-3′), L1CAM-specific siRNAs (sequence: 5′-GUGGAGUGAUCCUGGAUAUTT-3′,

5′-AUAUCCAGGAUCACUCCACTT-3′) and negative control siRNA (sequence: 5′-UUCUCCGAACGUGUCACGUTT-3′, 5′-ACGUGACACGUUCGGAGAATT-3′) were purchased from GenePharma (Shanghai, China). The pCMV3-entry (empty)vector and pCMV3-L1CAM-FLAG vector (with a FLAG tag at C-terminal) were purchased from Sino Biological (Beijing, China). The transfection of siRNAs and plasmid was performed using Lipofectamine 3000 kit (Invitrogen, Carlsbad, CA, USA) according to the manufacturer's protocol.

**Plasmid site-directed mutagenesis.** A mammalian expression pCMV3 vector containing the human L1CAM cDNA with a FLAG tag at C-terminal (pL1CAM-FLAG) was obtained from Sino Biological (Beijing, China). pL1CAM-FLAG plasmid was used as template to generate the pL1CAM (N979Q)-FLAG mutant by site-directed mutagenesis method. In brief, using the template-specific mutagenic primers (forward: 5'-CGAACTTCGGACACACCAGCTGACCGATCTCAGCC-3', reverse: 5'-GGCTGAGATCGGTCAGCTGGTGTGTCCGAAGTTCG-3'), the pL1CAM-FLAG plasmid was amplified by PCR with Phanta DNA polymerase (Vazyme, P501). The PCR products were digested with DpnI restriction enzyme, transformed into TOP10 competent cells, and the positive clones were picked up to confirm the corrected pL1CAM (N979Q)-FLAG mutant by DNA sequencing.
